# Beyond Inheritance: De novo Fast Motion Computation in Primate Visual Cortex

**DOI:** 10.64898/2025.12.02.691546

**Authors:** Keyan He, Lixuan Liu, Junxiang Luo, Yiliang Lu, Jiapeng Yin, Yingfan Liu, Wenheng Xie, Yizhejun Li, Xiaohong Li, Ian Max Andolina, Stewart Shipp, Hongbo Yu, Ye Wang, Dajun Xing, Niall McLoughlin, Wei Wang

**Affiliations:** Center for Excellence in Brain Science and Intelligence Technology, State Key Laboratory of Brain Cognition and Brain-inspired Intelligence Technology, Institute of Neuroscience, Chinese Academy of Sciences, Shanghai 200031, China; University of Chinese Academy of Sciences, Beijing 100049, China; School of Life Sciences, Fudan University, Shanghai 200433, China; State Key Laboratory of Media Convergence and Communication, Communication University of China, Beijing 100024, China; Neuroscience and Intelligent Media Institute, Communication University of China, Beijing 100024, China; State Key Laboratory of Cognitive Neuroscience and Learning & IDG/McGovern Institute for Brain Research, Beijing Normal University, Beijing 100875, China; Division of Pharmacy & Optometry, University of Bradford, UK

**Keywords:** sequential visuotopic activations, direction selectivity, velocity computation, cascaded spatiotemporal integration, dorsal visual pathway

## Abstract

Objects move through space and time, generating sequential visuotopic activations in all sighted animals leading to motion perception of velocity defined by direction and speed. Humans can effortlessly see motion with speeds ranging from 0.25 to 500°/s. However, direction-selective neurons in the primary visual cortex (V1)—from which all subsequent processing is presumed to derive— only encode directionality at low speeds. To resolve this paradox, we recorded neuronal responses to moving dots, gratings, and movies across the LGN, V1, MT and MST of the macaque motion pathway. Regardless of cell type and motion stimuli, V1 neurons lost direction selectivity at ∼29°/s while MT and MST neurons maintained it up to ∼82°/s and ∼183°/s, respectively. A cascaded spatiotemporal integration model reveals that at each cortex direction-selective neurons can generate velocity selectivity de novo, by integrating sequential visuotopic activations from preceding areas, irrespective of speed and directionality. By computing velocity anew, the primate brain effectively uses the cortical hierarchy itself to ‘shift gears’ to efficiently encode slow and fast motion. Thus, five visual areas from the retina into the brain’s processing hierarchy, external spatiotemporal information is being computed afresh, offering insights for motion processing in other species, modalities and machine vision.

## Introduction

Our ability to accurately perceive the direction and speed of moving objects in a fast-moving world is fundamental to our survival. For example, we can adjust our driving speed by changing gears according to the velocity of other road users (**Fig. 1a**). Velocity is a vector combining motion direction and speed, and is the core component of visual motion perception. It is derived physiologically across all sighted species from sequential retinotopic/visuotopic activations generated by moving stimuli (Bradley & Goyal, 2008; Dhande et al., 2015; Mauss et al., 2017). Motion processing is the ability to detect changes in the spatial position of an object over time (Mikami et al., 1986a, 1986b); in other words, direction cannot be defined without speed. Using a variety of moving stimuli, different studies have found that the range of optimal velocities encoded by direction-selective (DS) neurons varies significantly both within and across cortical motion areas of macaques, from 0.75 degree/second (°/s) to 4.47°/s in V1 (Dow, 1974; Orban et al., 1986; Priebe et al., 2006), 2°/s to 50°/s in MT (Zeki, 1974; Maunsell & Van Essen, 1983; Mikami et al., 1986a; Rodman & Albright, 1987; Lagae et al., 1993; Priebe et al., 2003), and 30°/s up to 160°/s in MST (Kawano et al., 1994; Orban et al., 1995; Miura et al., 2014). The variation of velocity at each area also depends on the eccentricity of the DS neurons recorded, with neurons preferring much slow velocity in the central visual field. Regardless, DS response peaks around their respective optimal speeds for V1 and MT. In human subjects, the optimal velocity was found to be around 3.7°/s for V1 and ranged from 7°/s to 30°/s for MT (Chawla et al., 1998).

**Fig. 1.**
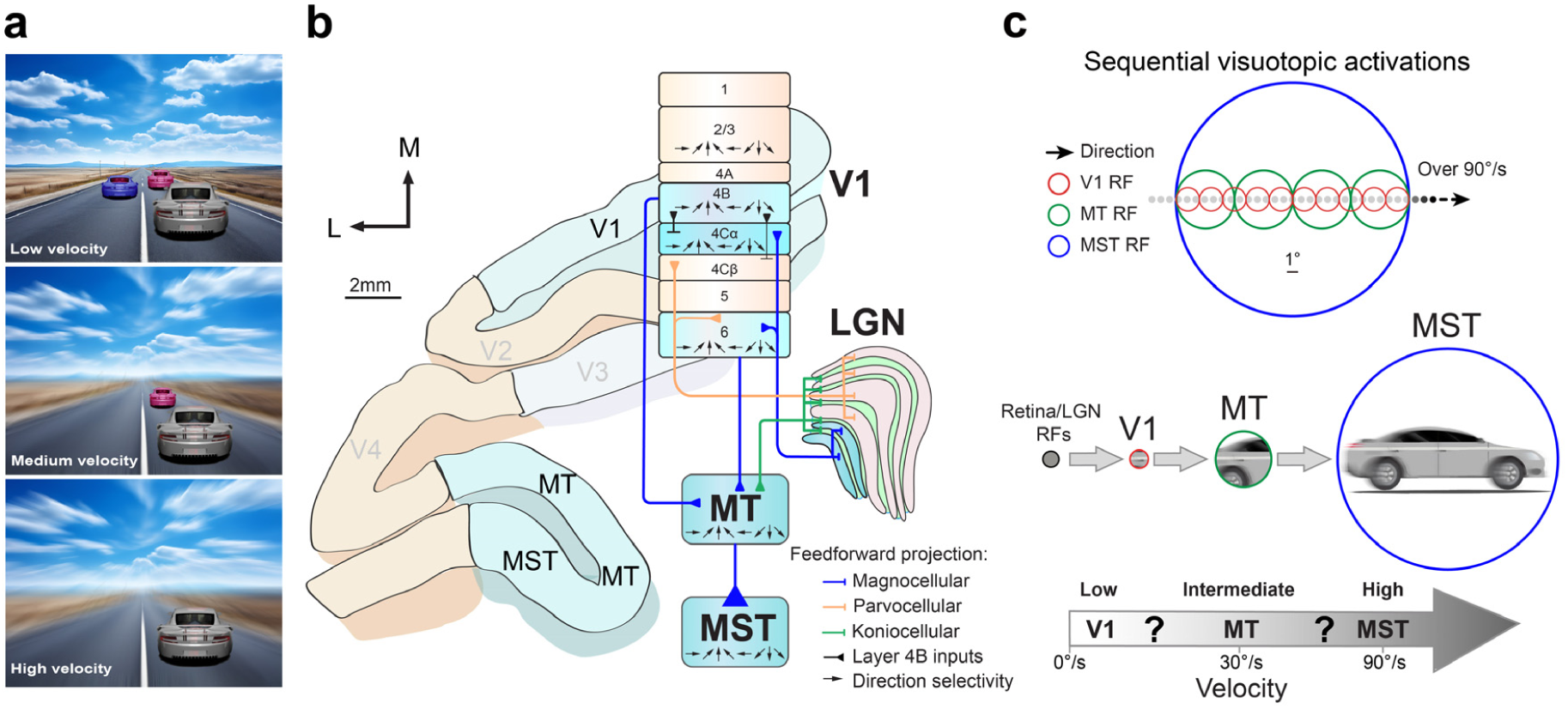
Overview of the psychology, anatomy, and physiology for primate velocity computation. **a.** Velocity perception is critical for humans in a dynamic world. **b.** Anatomical projections in primate dorsal visual pathway. DS neurons generated from major ascending projections from non-DS neurons in lateral geniculate nucleus (LGN) to the primary visual cortex (V1), the middle temporal (MT) and the medial superior temporal (MST) areas within the primate dorsal visual stream, processing visuotopic motion signals. **c.** Superimposed receptive fields (RFs) of DS neurons in V1 and MT, but not in MST, exhibit visuotopic organization. RF size is dramatically increased along the visual hierarchy of V1, MT, and MST, correlating with the ability to deal with faster motion, but how, particularly for the transforms of intermediate and high velocities after V1 loses its direction selectivity at its cutoff velocity remains unknown.

**Fig. 2.**
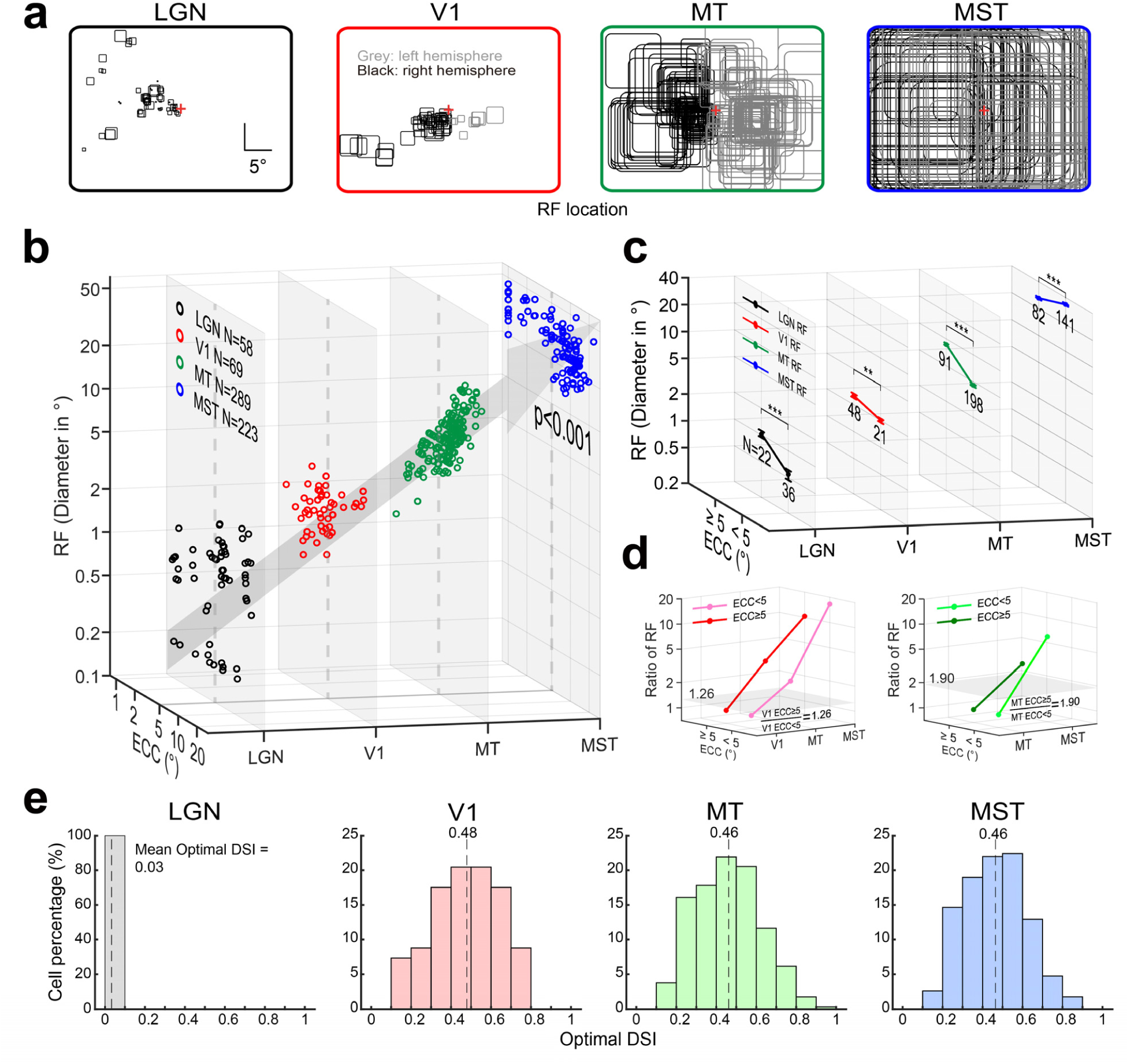
Receptive field properties and the optimal DSI of recorded neurons along the hierarchy. **a.** The RF mappings of neurons across four visual areas along the hierarchy. The black and gray RFs illustrate the right and left hemispheres, respectively. **b.** The distribution of RF sizes in diameters along eccentricity in each brain area. The RF diameters of LGN, V1, MT and MST are consistent with previous reports. The RF diameter: 0.83±0.07°, 2.11±0.09°, 7.35±0.20° and 28.85±0.49°, for LGN, V1, MT, and MST, respectively (Kruskal-Wallis test, p<0.001). **c.** The comparison of RF diameters between above and below 5° eccentricities across LGN, V1, MT and MST. Single, dual and triple stars represent statistical significance with p<0.05, p<0.01, p<0.001, respectively, in these and subsequent charts. **d.** The changes of the RF diameters across cortices relative to V1 and MT neurons, respectively. The horizontal gray surfaces indicate the ratio levels of RF diameters for neurons above and below 5° eccentricities. **e.** The distribution of DSI for all recorded neurons across the four areas along the visual hierarchy.

However, humans can perceive an enormous range of velocities, with speeds ranging from 0.25 to 500°/s being used experimentally (Orban et al., 1984; Kawakami et al., 2002). It remains unknown how this wide range of velocities are computed and transformed by DS neurons with huge differences of receptive field (RF) sizes across cortices of the primate dorsal visual motion stream (Born & Bradley, 2005; Bradley & Goyal, 2008; Nassi & Callaway, 2009) (**Fig. 1bc**). Previous studies using simple dots, sine-wave gratings and plaids to map direction and speed selectivity have predominantly focused on a limited range of optimal speeds in one or at most two cortical regions (Dow, 1974; Zeki, 1974; Maunsell & Van Essen, 1983; Mikami et al., 1986a, 1986b; Orban et al., 1986; Rodman & Albright, 1987; Lagae et al., 1993; Kawano et al., 1994; Orban et al., 1995; Chawla et al., 1998; Priebe et al., 2003; Priebe et al., 2006; Miura et al., 2014). This helps explain why the key question— how slow and fast velocities across the full spectrum are encoded and transformed in the dorsal visual pathway—has, until now, remained unaddressed.

Based on early anatomical and physiological findings that inputs to MT derive specifically from V1 layers 4B and 6 in which DS neurons are concentrated (Shipp & Zeki, 1989; J. A. Movshon & Newsome, 1996; Nassi & Callaway, 2009) (**Fig. 1b**), it is generally held that higher cortical areas such as MT inherits their direction selectivity from V1 (Born & Bradley, 2005; Churchland et al., 2005; Priebe et al., 2006). However, DS neurons in V1 only encode directionality at much low speeds. The RFs of DS neurons in V1 are the smallest of all cortices in the hierarchy, and as such, objects moving above a certain speed flash by as they only appear for a very short period of time within the RFs of V1 DS neurons. Therefore, V1 DS neurons only respond with transient nondirectional visuotopic activations at higher velocities (**Fig. 1c**). This raises a critical question: how do neurons in higher cortical areas maintain direction selectivity once V1 loses it at high speeds? One possibility is that a small subset of V1 neurons retains selectivity at higher velocities, providing a “labeled-line” that MT can sample as speed increases. This would require an unknown mechanism so that small receptive fields of V1 DS cells integrate motion beyond their boundaries, and MT switching between labeled lines to preserve selectivity. A more plausible explanation is that larger receptive fields naturally extend the time a moving stimulus remains within them. Thus, as receptive fields expand along the visual hierarchy, DS neurons can process progressively faster velocities in a stepwise fashion. This parallels the idea that the larger receptive fields of MT DS neurons help resolve the ‘aperture problem’ that constrains V1 DS neurons with smaller RFs (Pack & Born, 2001; Bradley & Goyal, 2008). The fine substructure of inputs from V1 DS cells can be measured within the larger RFs of MT cells (M. S. Livingstone et al., 2001; M. S. Livingstone & Conway, 2003), demonstrating that MT DS neurons spatiotemporally pool spike inputs from multiple presynaptic neurons across space (Hubel & Wiesel, 1962; Reid & Alonso, 1995; M. S. Livingstone, 1998; De Valois et al., 2000; M. S. Livingstone et al., 2001; M. S. Livingstone & Conway, 2003) in a way similar to V1 DS cells integrating their LGN inputs spatially (Hubel & Wiesel, 1962; J. A. Movshon et al., 1978; Bradley & Goyal, 2008). Similar to macaque V1 layer 4Cα DS simple cells, direction selectivity of mouse V1 simple cells can also emerge anew independent of the direction selectivity of the presynaptic neurons (Lien & Scanziani, 2018; Rossi et al., 2020). Finally, evidence from patients with V1 lesions demonstrates that MT+ retains motion abilities using residual inputs alone (Rodman & Albright, 1989; Rosa et al., 2000; Azzopardi et al., 2003; Alexander & Cowey, 2009; Huxlin et al., 2009; Das et al., 2014; Ajina et al., 2015).

Supporting this idea is perhaps the most fundamental property of the visual system in all species: retinotopic/visuotopic organization. Nearby locations in the retina, corresponding to nearby locations in the visual world, form mosaic-like maps to corresponding cortical neurons (Tootell et al., 1982). Sequential retinotopic activations, generated by a moving object, were used to study selectivity to motion direction in the retina of insects and rabbits, and led to the development of the Hassenstein- Reichardt and Barlow-Levick motion detector models more than sixty-five years ago (Hassenstein & Reichardt, 1956; Barlow & Levick, 1965; Mauss et al., 2017). More recently, these sequential visuotopic activations in V1 have been captured mathematically by a spatiotemporal energy model of direction selectivity (Adelson & Bergen, 1985; Simoncelli & Heeger, 1998; Bradley & Goyal, 2008), mapped experimentally by optical imaging (Jancke et al., 2004; Lu et al., 2017), and also serving the development of cortical visual prosthesis (Beauchamp et al., 2020; X. Chen et al., 2020). We propose that this fundamental property of sequential visuotopic activations, modelled by spatiotemporal motion energy detectors, forms the basis of how not only V1 but also how higher visual areas (MT and MST) generate velocity selectivity. At higher velocities, when V1 neurons lose their direction selectivity, they are still retinotopically activated in a manner similar to retinal and LGN neurons. An object moving at high velocity will thus cause sequential activations within the multiple visuotopic maps of the LGN, V1, and higher visual areas that are retinotopically organized. By employing the same principles used for generating V1 direction selectivity from sequential visuotopic inputs of LGN neurons, we propose that MT and MST can integrate spatiotemporal inputs of preceding cortical areas to generate velocity selectivity *de novo*. A separate direct projection also exists from the LGN to MT (Sincich et al., 2004; Nassi & Callaway, 2009), which may also contribute to the construction of DS neurons’ spatiotemporal RFs in MT. Other inputs to MT include a disynaptic pathway from superior colliculus and pulvinar related to eye movements (Lyon et al., 2010), and from V2/V3 for binocular disparity rather than DS responses (Ponce et al., 2008).

In this study, we were able to directly address the above fundamental question using MRI guided extracellular single-cell recordings of the neuronal responses across four successive regions: LGN, V1, MT and MST in the dorsal visual stream of the same awake macaques to moving stimuli composed of dots, sine-wave gratings and natural images across a wide range of velocities. Both optimal and cutoff velocities were measured for DS neurons within V1, MT and MST. To model our findings we modified Simoncelli and Heeger 1998’s model of V1 direction selectivity to incorporate inputs from both the LGN and V1 DS cells to generate velocity selective responses in MT (Adelson & Bergen, 1985; Simoncelli & Heeger, 1998; Nassi & Callaway, 2009) and applied the same model to the outputs of MT to generate velocity responses in MST. We propose that, regardless of cell type or motion stimulus, the same spatiotemporal integration mechanism that generates velocity selectivity within the eyes of insects and rabbits can also explain our findings across a wide range of velocities in natural environments. In doing so we evoke the law of parsimony, or Occam’s razor, whereby the simplest explanation is often the best one, a principle that goes back at least as far as Aristotle in 300 BC, who wrote "Nature operates in the shortest way possible".

## Results

MRI guided extracellular recordings of four successive visual areas of LGN, V1, MT and MST along the dorsal visual pathway were accomplished in four behaving macaques performing a fixation task (**Supplementary Fig. 1ab**). The spatial frequency (SF) distribution of all stimuli used are as presented (**Supplementary Fig. 1c**). Movies consisting of natural images of trees and leaves were used as an additional stimulus to test subsamples of MT and MST DS neurons. A total of 58 non-directional LGN neurons, and 69 V1, 289 MT, 223 MST DS neurons were recorded, respectively, in response to moving fields of random dots. The RF dimensions of DS neurons increase significantly along the visual hierarchy, and the eccentricities of all neurons recorded in LGN, V1, MT, and MST ranged from 1° up to 25° (**Fig. 2a-d**). Some neurons in MST have huge RFs that were larger than the screen and so we set their eccentricities to be centered on 0°. Only neurons with a maximum direction- selective index (DSI) higher than 0.1 were recorded in V1, MT and MST. The optimal DSI ranged from 0.1 to 0.9 across populations, and the average DSI is very similar among the three cortices (**Fig. 2e**). Some of the DS neurons were also examined with drifting sine-wave gratings for the classification of simple and complex cells (Skottun et al., 1991), and component and pattern cells (J. Movshon et al., 1985; Rodman & Albright, 1989; Khawaja et al., 2009).

### Velocity Representation across Cortices

**Fig. 3a** illustrates responses from four example neurons recorded in the LGN, V1, MT and MST to fields of random dots moving at various velocities. Neuronal responses were fitted with a Gaussian function to find the optimal responses in each velocity tuning curve (*methods section*). Across the populations recorded, LGN cells did not show any direction selectivity, giving strong responses at all velocities tested, consistent with previous studies (Wiesel & Hubel, 1966; Derrington & Lennie, 1984). However, DS neurons in V1, MT and MST with similar direction preferences display huge differences in their optimal and cutoff velocities. Optimal velocity is defined as the velocity at which DS neurons exhibit their maximum spike rates, while the cutoff velocity is defined as the velocity at which the DSI of a DS neuron drops below 0.1. The optimal and cutoff velocities for the example V1, MT and MST neurons are 11°/s, 28°/s, 59°/s and 35°/s, 73°/s, >200°/s, respectively. The responses from further examples of different groups of neurons are presented in **Supplementary Figs.2-3**. Both optimal and cutoff velocities are drastically increased along the hierarchy in each of all four monkeys. Population results of four monkeys are presented, for direct comparisons of the optimal and cutoff velocities across the three cortices (**Fig. 3bc**). V1 is sharply tuned to lower velocities whereas MT and MST are more broadly tuned and cover a much wider range of medium to higher velocities. The bandwidths of MT and MST velocity tuning curves are also significantly wider than those in V1. The optimal and cutoff velocities of V1, MT and MST populations are 9°/s, 25°/s, 48°/s, and 29°/s, 82°/s, 183°/s, respectively. Both optimal and cutoff velocities are drastically increased along the hierarchy. Importantly, V1 accurately encodes low velocities, however, by its cutoff velocity (29°/s) V1 has lost its DS responses whereas MT neurons retain strong DS responses until 82°/s (its cutoff velocity). The cutoff velocities of V1 and MT DS neurons are consistent with previous reports for the study of spatial limits of DS responses in V1 and MT (Mikami et al., 1986b; Churchland et al., 2005). The cutoff velocity for MST DS neurons is the highest of the areas investigated (> 150°/s), well above the cutoff velocities at which both V1 and MT successively lose their direction selectivity (**Fig. 3a-c**). In addition, the optimal firing-rate velocities and the optimal DSI velocities are highly compatible for V1, MT and MST (**Fig. 3d**). Importantly, the average responses to moving dots beyond the cutoff velocity for each area is more than twice their spontaneous firing rates (without a stimulus present, **Fig. 3e**).

We further tested samples of DS neurons along the visual hierarchy in response to drifting sine- wave gratings. The optimal velocities of drifting sine-wave gratings can be computed as the ratio of optimal temporal frequency (TF) over optimal SFs (Perrone & Thiele, 2001; Priebe et al., 2003; Priebe et al., 2006). Simple and complex cells in V1 can be classified based on their Fourier (linear) modulations of visual responses to drifting sine-wave gratings (method section). With the same criterion, we grouped simple and complex types of DS neurons in both MT and MST for a direct comparison with V1 (**Fig.4 and Supplementary Fig.2**). It has been reported that there is no statistical difference between optimal velocities or bandwidths of V1 simple and complex cells (Priebe et al., 2006). This is confirmed here not only in V1, but also in MT and MST as well as in their responses to either drifting sine-wave gratings or moving dots (**Supplementary Fig.4**). This also applies to the comparison of cutoff velocities and bandwidths within each cortex between simple and complex cells (**Fig.4 e-g**). Thus, regardless of cell types, the optimal and the cutoff velocities, and the bandwidths of DS neurons increased massively across the three cortices along the visual hierarchy.

**Fig. 3.**
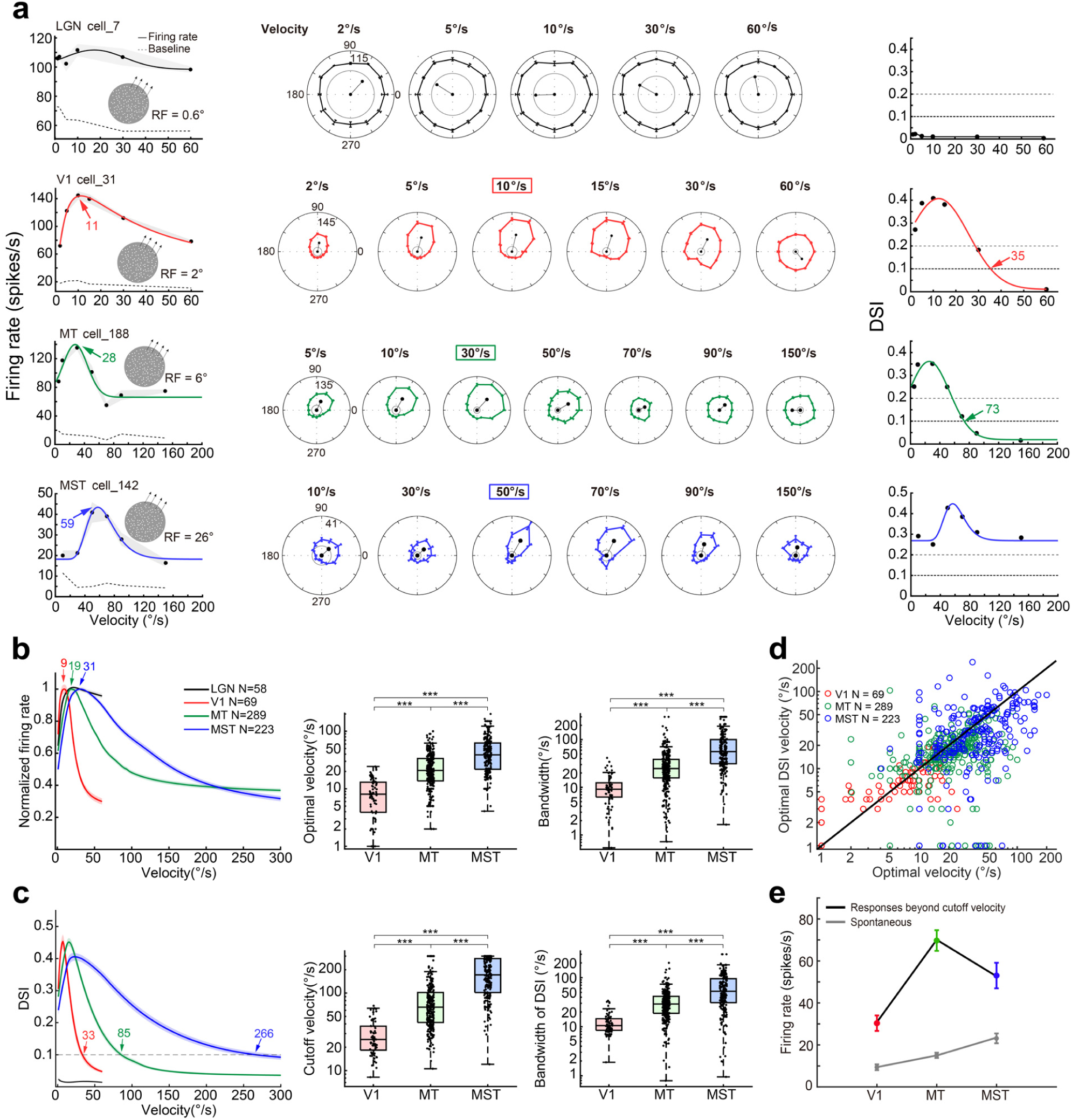
The transformation of velocity from low to high along the dorsal visual stream. **a.** Four example neurons recorded from LGN, V1, MT and MST show distinctive responses to dot motion at various velocities. Left column: velocity tuning responses. The dotted lines indicate spontaneous activity when no stimulus presented. Middle panel: direction tuning curves in circular plots. The colored boxes indicate the optimal velocities of each neuron, respectively. Right column: DSI velocity tuning curves. The colored digits with arrow points indicate the optimal and cutoff velocities, respectively. The shade and error bar indicate the SEM. **b-c.** The optimal and cutoff velocity comparison across three areas. The left panel exhibits the average tuning curve of individual DS neurons in V1, MT and MST. The colour digit above each curve indicates the optimal and cutoff velocities of the averaged curves. The middle and right panels present the boxplots of optimal and cutoff velocities along with bandwidths of populations in three areas. Single, dual and triple stars represent statistical significance with p<0.05, p<0.01, p<0.001, in these and subsequent charts. **d.** The velocity comparison between optimal DSI and optimal responses. **e.** The population comparison between average responses after cutoff velocities and spontaneous responses across V1, MT and MST.

**Fig. 4.**
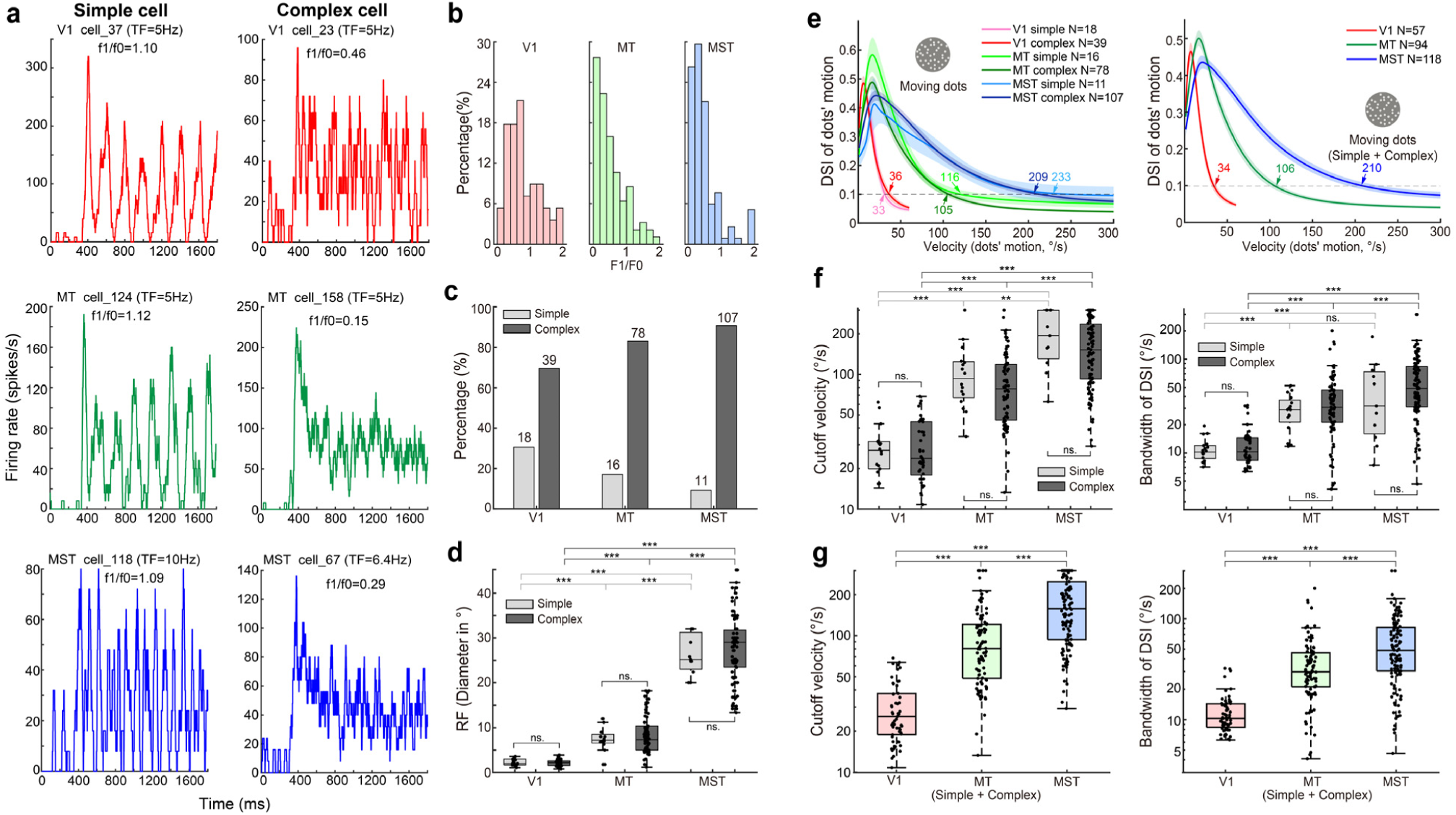
The classification and examples of simple and complex cells, and the DSI tuning responses to random dot motion in V1, MT and MST along the visual pathway. **a.** The PSTH of simple and complex cells in response to drifting sine-wave gratings. **b.** F1/F0 distribution of simple and complex cells along the hierarchy. **c.** The proportion of simple and complex cells along the hierarchy. **d.** The comparison of RFs between simple and complex cells within and across cortices along the hierarchy. The RF diameters of simple cells: 2.25±0.17°, 7.39±0.74°, and 27.38±2.17°, for V1, MT, and MST, respectively. The RF diameters of complex cells: 2.16±0.11°, 8.37±0.49° and 27.75±0.71°, for V1, MT, and MST, respectively. The ns. here and elsewhere represents no statistical significance with p≥0.05. **e.** The population DSI tuning curves of simple and complex cells to dots’ motion field across V1, MT and MST, and the DSI tuning curves without separating simple and complex cells along the hierarchy. **f.** The comparison of cutoff velocities and bandwidths of simple and complex cells along the hierarchy. The cutoff velocity of simple cells: 29.11±3.13°/s, 108.36±16.14°/s and 201.95±26.75°/s for V1, MT and MST, respectively. The cutoff velocity of complex cells: 29.47±2.50°/s, 95.67±7.46°/s and 166.89±9.55°/s for V1, MT and MST, respectively. The bandwidth of simple cells: 10.98±0.77°/s, 29.09±3.06°/s and 51.80±14.78°/s for V1, MT and MST, respectively. The bandwidth of complex cells: 12.61±1.05°/s, 39.75±3.83°/s and 62.73±4.78°/s for V1, MT and MST, respectively. **g.** The comparison of the cutoff velocity and bandwidth of populations across V1, MT and MST.

In addition, we also tested more than half the population of V1, MT and MST neurons we recorded from using component and pattern motion stimuli (J. Movshon et al., 1985; Khawaja et al., 2009). We classified and grouped our DS neurons into component, pattern and unclassified groups (**Fig.5ab and Supplementary Fig.3**). The distribution of the three neuron groups within each cortex of the hierarchy is consistent with previous studies (Khawaja et al., 2009) (**Fig.5ab**). Importantly, velocity tuning responses for all three cell groups are presented and compared (**Fig.5c-f**). Regardless of their differing roles in determining object motion direction, there is no statistical difference for both their optimal and cutoff velocities within each cortex of V1, MT and MST, respectively. However, the optimal and cutoff velocities of the three different groups demonstrate the familiar stepwise increase along the visual hierarchy. While random dots are a popular stimulus for investigating visual motion, by virtue of mimicking natural scenes through their content of multiple SFs (**Supplementary Fig.1c**), we also confirmed these findings using a naturalistic movie containing moving leaves and jungle scenes (**Supplementary Fig.5**).

**Fig. 5.**
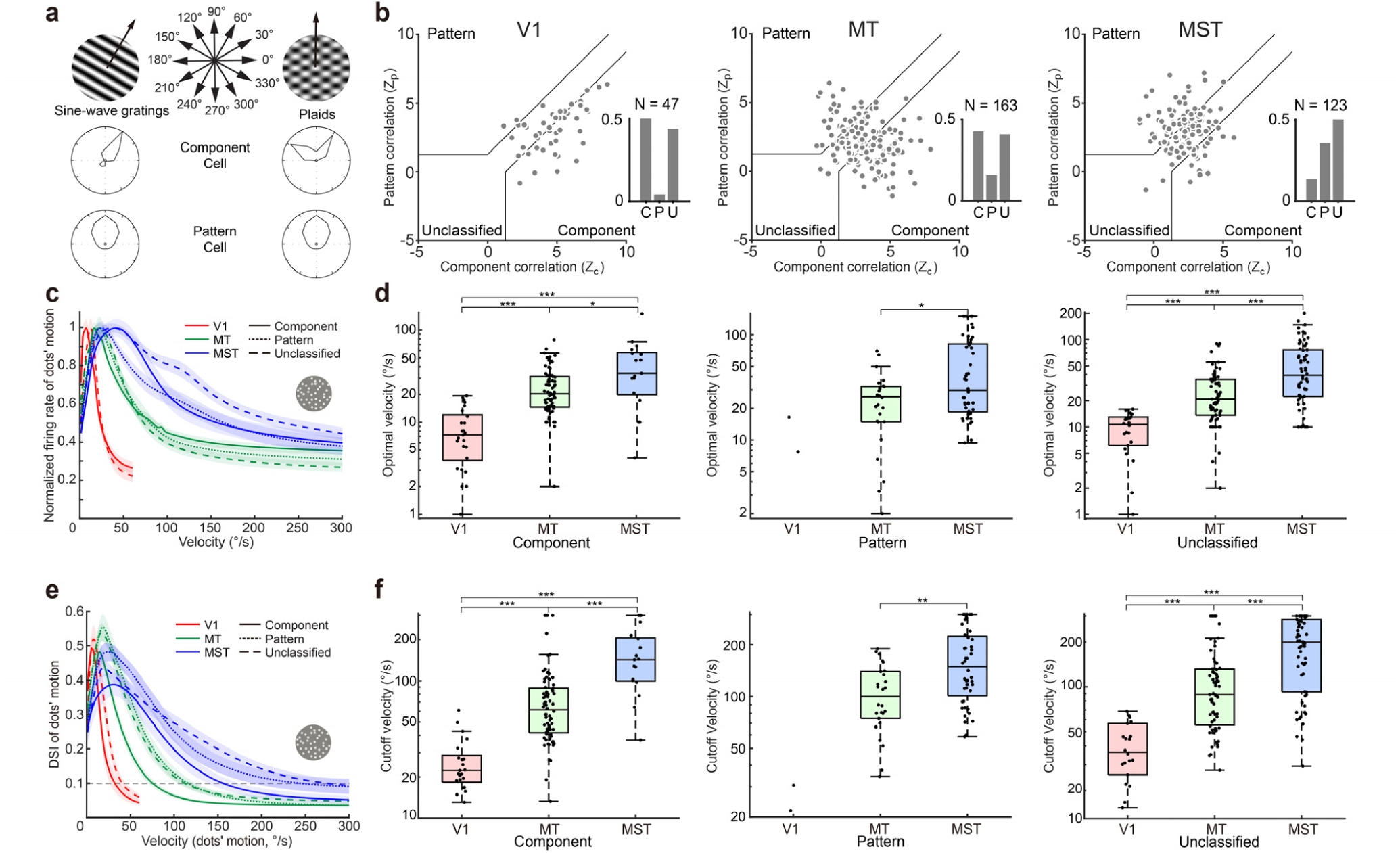
The population velocity tuning responses of component, pattern and unclassified cells to moving fields of random dots along the hierarchy. **a.** The classification of component and pattern cells. **b.** The proportion and distribution of component, pattern and unclassified cells along the hierarchy. **c.** The velocity tuning responses of component, pattern and unclassified cells across V1, MT and MST. **d.** The comparison of optimal velocities among three types of cell groups along the hierarchy. The optimal velocity of component cells: 8.43±1.14°/s, 25.05±1.75°/s and 43.81±8.38°/s for V1, MT and MST, respectively. The optimal velocity of pattern cells: 25.68±3.29°/s and 52.37±6.69°/s for MT and MST, respectively. Note that only two pattern cells were recorded in V1, which were presented but not included for statistic comparison with those in MT and MST. The optimal velocity of unclassified cells: 9.78±1.00°/s, 26.62±2.34°/s and 54.06±5.38°/s for V1, MT and MST, respectively. **e.** The DSI tuning responses of component, pattern and unclassified cells across V1, MT and MST. **f.** The comparison of cutoff velocities among three types of cell groups along the hierarchy. The cutoff velocity of component cells: 25.51±2.41°/s, 77.18±7.06°/s and 154.69±19.13°/s for V1, MT and MST, respectively. The cutoff velocity of pattern cells: 105.83±8.71°/s and 166.32±11.96°/s for MT and MST, respectively. The cutoff velocity of unclassified cells: 38.57±3.56°/s, 113.50±9.29°/s and 191.36±11.52°/s for V1, MT and MST, respectively.

In summary, we found that both the optimal and cutoff velocities for DS neurons increased significantly across V1, MT and MST regardless of how we classified DS neurons. This change in optimal and cutoff velocities between cortical areas defies the standard model that DS in higher cortical areas is exclusively inherited from DS neurons in V1. How do neurons in MT and MST respond in a direction-selective manner to stimuli moving at a velocity faster than the cutoff velocity of V1?

### Velocity transformation from slow to fast

Drifting sine-wave gratings are commonly used to study spatiotemporal properties of DS neurons’ RFs in macaque V1 and MT. Within our sample of DS neurons, the average optimal SFs decreased dramatically along the hierarchy of V1, MT and MST (**Supplementary Fig.6a-c**). Clearly, the optimal SFs of DS neurons are inversely correlated with their RF diameters along the hierarchy. DS neurons in MT and MST encode high velocity within their greatly enlarged RFs, but at the expense of visual acuity when compared to V1. In contrast, the average optimal TFs of DS neurons across the cortices only weakly correlated with the increase in RF diameters, although V1 optimal TF is still significantly lower than those in MT and MST (**Supplementary Fig.6d-f**). This is consistent with previous findings (Wang & Movshon, 2016). There was no statistical difference between the optimal TFs of neurons in MT and MST. RF sizes change not only between cortical areas but also with eccentricity within individual cortical regions. To investigate this further, we plotted the optimal SFs and TFs of DS neurons against their eccentricity within V1, MT and MST (**Supplementary Fig.6g-l**). Within our sample of V1, MT and MST neurons, there is only a modest effect of eccentricity upon optimal SFs and TFs, partly due to the uneven distribution of recorded eccentricities. Compared to this limited effect of eccentricity, the changes of both SFs and TFs across cortices along the hierarchy are much more pronounced. More importantly, the increase in both optimal and cutoff velocities of DS neurons from V1 to MT and MST correlates with the huge increase in their RF diameters across these regions and is inversely correlated with the change in their optimal SFs (**Fig.6ab**). Optimal TFs along the hierarchy are much less relevant when compared with optimal SFs (**Fig.6c**).

**Fig. 6.**
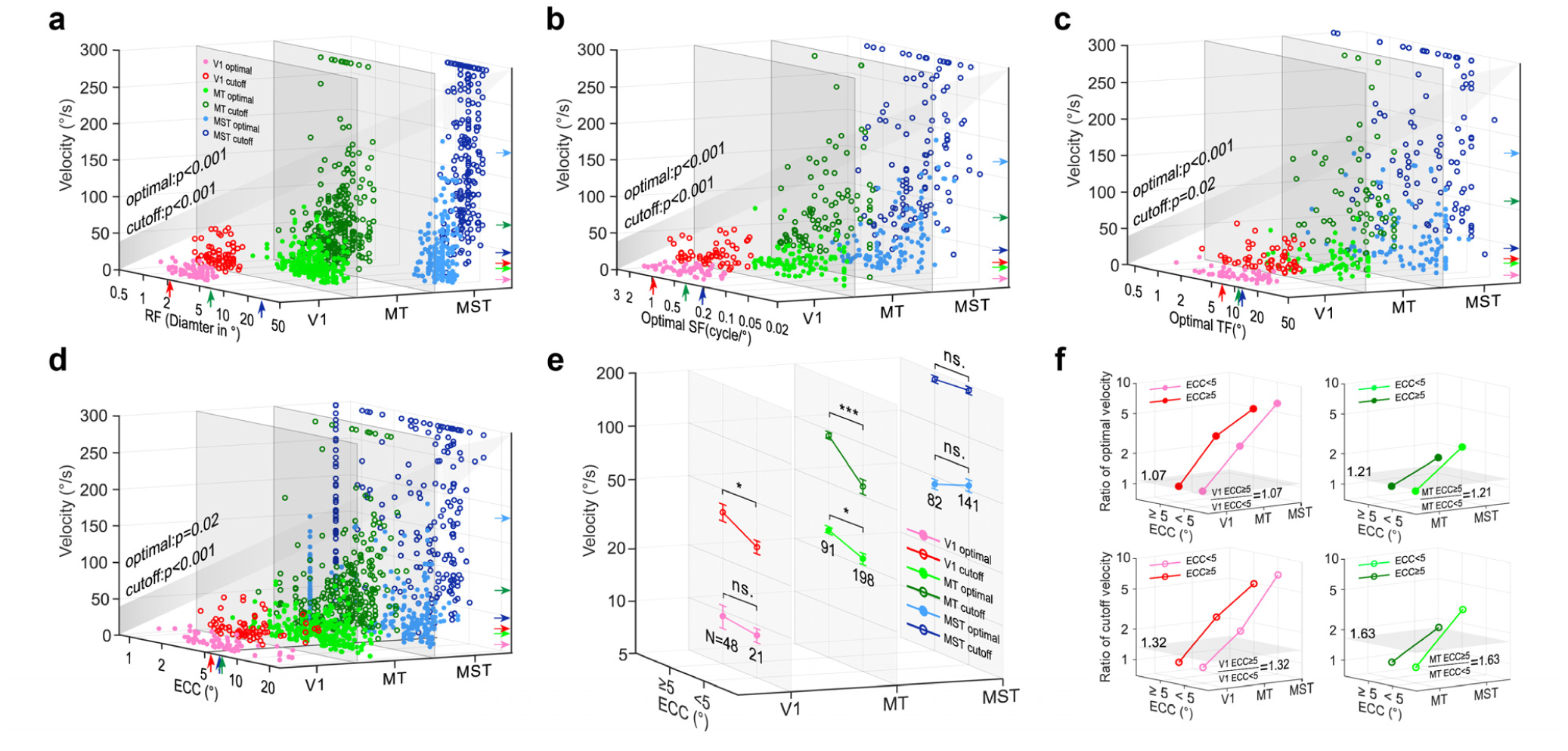
The velocity transformation governed by DS neuron spatial characteristics across visual hierarchy and eccentricity (ECC). **a-d.** The optimal and cutoff velocities are plotted against RFs’ diameters, SFs, TFs, and eccentricity, respectively, along the visual hierarchy. The color filled dots and hollow circles represent optimal and cutoff velocities, respectively. The p-values illustrate the significance of correlations. In panel A to D, the vertical color arrows represent the average values of RF, optimal SF, optimal TF, and ECC of each area, respectively. The horizontal color arrows indicate the average values of optimal and cutoff velocities for each area, respectively. **e.** The comparison of optimal and cutoff velocities below and above 5° eccentricity within and across V1, MT and MST. **f.** The relative changes of the optimal and cutoff velocities across cortices after normalizing to V1 and MT, respectively. The gray horizontal surfaces indicate the ratio levels of the optimal and cutoff velocities for V1 and MT neurons, respectively, above and below 5° eccentricity.

Previous studies have found that cutoff velocities (defined by decreases of firing rates) increase with eccentricity for DS neurons of areas 17 and 18 (Orban et al., 1986). We investigated eccentricity effects on both optimal and cutoff velocities of DS neurons across V1, MT and MST cortices. Overall, the optimal and cutoff velocities increased across eccentricity along the hierarchy (**Fig.6d**). Specifically, cutoff velocities are significantly increased across the eccentricity for DS neurons in retinotopic organized V1 and MT, but not MST (**Fig.6e**). Optimal velocity increases significantly for MT but not for V1 DS neurons, likely because much of our sample of V1 cells is clustered within 1∼5° eccentricity. Regardless, the impact of eccentricity is much less for both the optimal and cutoff velocities than that of changing cortical areas (**Fig.6f**). Overall, our results demonstrate that the encoding of velocity from low to high is governed mainly by variation of the spatial characteristics of DS neurons between cortices in the visual hierarchy, i.e. the larger the RF size of DS neurons along the dorsal motion pathway, the faster the velocity they encode.

### De Novo Cortical Velocity Selectivity

All models of direction selectivity comprise computation of a time delay between adjacent inputs in the retina (Hassenstein & Reichardt, 1956; Barlow & Levick, 1965). In primates, moving objects initiate neuronal spikes from the retina and subsequently from the LGN before feeding into V1 (**Figs.1b and 7a**). DS responses in V1 simple cells can be successfully explained in both experiments and computational models as a result of spatiotemporal integration of LGN inputs (Adelson & Bergen, 1985; Reid et al., 1987; Jagadeesh et al., 1997; De Valois et al., 2000). **Fig.7a** presents a diagram depicting the simplest feedforward model of DS responses generated by integrating non-DS LGN neuronal responses to V1, and then from V1 to MT and up to MST along the visual hierarchy. Sequential retinotopic/visuotopic activations are represented across all regions regardless of the velocity of the stimulus. In other words, spatiotemporal feedforward inputs are not only responsible for the generation of DS responses in V1 from non-direction-selective LGN inputs at low speed (DeAngelis et al., 1993; De Valois et al., 2000; Baker & Bair, 2012; Chariker et al., 2022), but these same visuotopic activations could also be responsible for the generation of DS responses in MT and MST at high speeds above the V1 cutoff velocity where V1 loses its directionality (**Fig.1c and 3e**). We propose that this cascaded integration of spatiotemporal information across cortices is the fundamental principle by which velocity selectivity can be generated at each motion processing stage (**Supplementary Fig.7**).

**Fig. 7.**
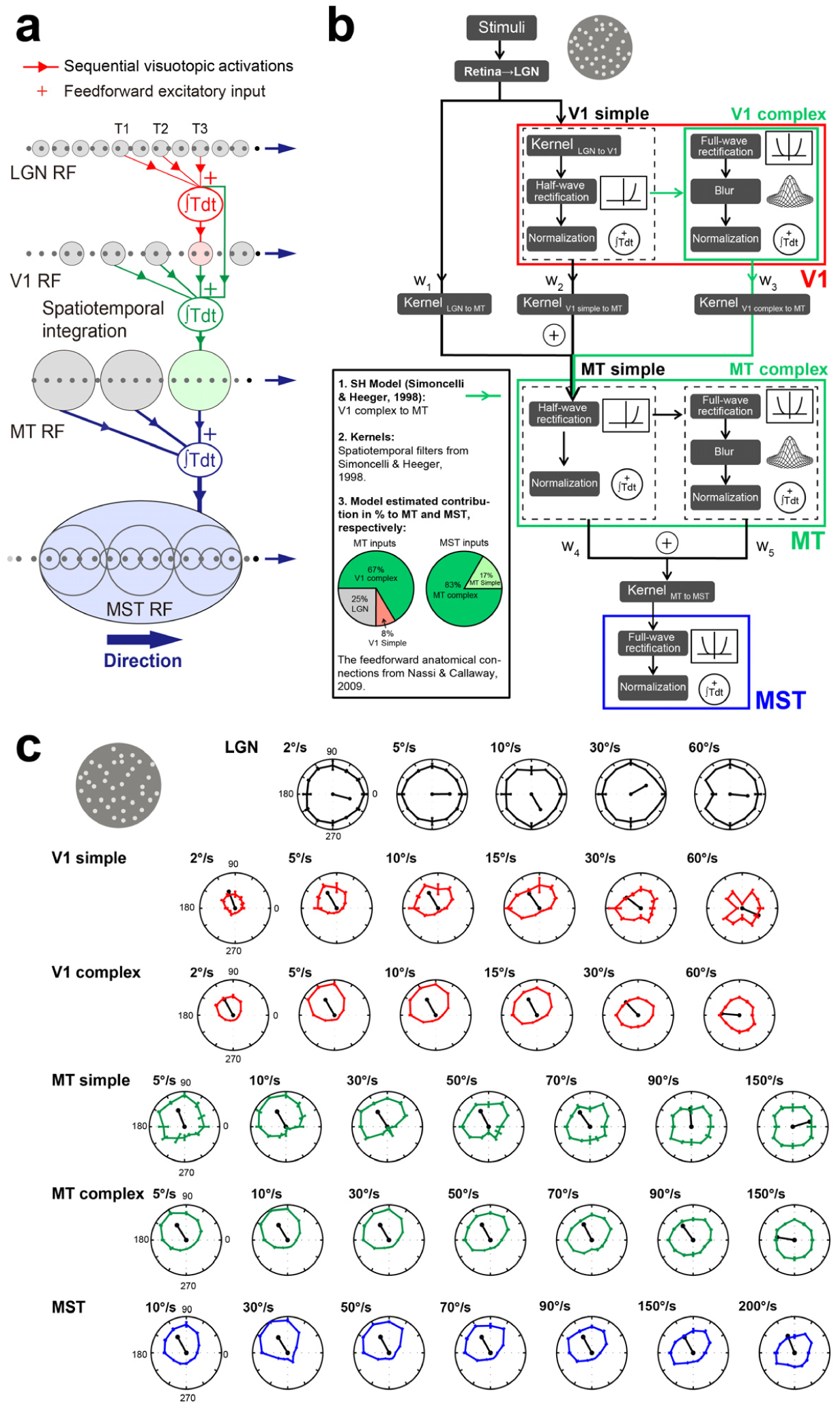
The simulation of the cascaded spatiotemporal integration (CSTI) model for velocity responses across LGN, V1, MT and MST along the dorsal visual pathway. **a.** A cascaded diagram of the Hassenstein-Reichardt motion detector across primate LGN, V1, MT and MST. Motion processing is the ability to discriminate the spatial position change (△x) of a moving object over time (△t). The diagram illustrates the simplest feedforward spatiotemporal inputs of visuotopic activations across cortices along the hierarchy. **b.** The structure and computational operations of cascaded spatiotemporal integration model. **c.** The simulation results of cascaded spatiotemporal integration model. The direction circular tuning curves at various speeds across LGN, V1, MT and MST are reproduced by the cascaded spatiotemporal integration model.

To confirm the above hypothesis, we adapted Simoncelli and Heeger’s model (Simoncelli & Heeger, 1998; Bradley & Goyal, 2008) by taking the same spatiotemporal kernels (Freeman & Adelson, 1991; Simoncelli & Heeger, 1998) to construct a cascaded spatiotemporal integration (CSTI) model, in order to simulate velocity computation across V1, MT and MST of the dorsal visual pathway (**Fig.7b and Supplementary Fig.7**). The V1 DS responses are simulated conventionally by integrating the outputs of basic spatial-temporal filters (**Supplementary Fig.7a**), which are based on the known neuronal activity of LGN and thalamo-cortical synaptic processes (Reid et al., 1987; DeAngelis et al., 1993; Jagadeesh et al., 1997; Simoncelli & Heeger, 1998). The fitting constraints for the RF sizes were set based on our experimental results (**Fig.2b**). The connection weights were initially set according to the existing literatures (Freeman & Adelson, 1991; Simoncelli & Heeger, 1998; Sincich et al., 2004; Bradley & Goyal, 2008; Nassi & Callaway, 2009) and adjusted to best fit the experiment data. In this model, neurons in downstream higher cortical regions are driven by maximum responses of preceding neurons when stimuli are presented at their preferred motion directions (**Supplementary Fig.7b**). For the simulation of MT DS responses, apart from the inputs from V1, we added a model component that captured the direct connection between the LGN and area MT (Sincich et al., 2004; Nassi & Callaway, 2009). These LGN inputs bypass V1 and may directly contribute to the construction of simple-cell type RFs of some MT neurons (M. S. Livingstone et al., 2001). In addition, a direct contribution from V1 simple cells to MT is also added into the model (Sincich et al., 2004; Nassi & Callaway, 2009). Within the model, the connection strengths from the LGN to V1 and to MT and from MT to MST were set to best simulate our experimental findings without considering any modulation of recurrent connections. We found that inputs from both the LGN and V1 were necessary to model the MT DS responses reported. This was further confirmed by our cascaded spatiotemporal integration model simulations with and without LGN and V1 inputs, respectively (**Supplementary Fig.8**). By varying the model parameters to reflect the spatial scale changes between cortices, the tuning responses of LGN, V1, MT and MST cells to moving dots over a wide range of velocities were generated, from which the optimal and cutoff velocities of DS neurons along the hierarchy are closely reproduced (**Figs.7c and 8a**).

Taken together and based on our current knowledge of anatomical projections between cortices along the hierarchy (Shipp & Zeki, 1989; Nassi & Callaway, 2009), the cascaded spatiotemporal integration model can reproduce the reported velocity responses for each cortex. The cascaded spatiotemporal integration model is capable of generating DS responses across V1, MT and MST at both low speeds (<30°/s), and at higher speeds (>30°/s) when V1 becomes insensitive to direction but DS neurons in MT and MST remain directionally sensitive (**Figs.7 and 8a**). In short, all of our experimental and simulation results demonstrate that the RFs of DS neurons in all cortical regions essentially serve as simple and efficient spatiotemporal motion energy detectors and in doing so accommodate a wide range of velocities in natural environments (**Fig.8b and Supplementary Fig.7bc**).

## Discussion

In natural environments, all sighted animals have to process a wide range of slow- and fast-moving objects travelling at different velocities. The visual brains in mammals and particularly primates have evolved hierarchically organized cortices dedicated to the visual tasks that nature imposes. Retinotopic organization, coupled with the orientation and/or direction selectivity of visual neurons, are the most fundamental building blocks of early visual processing. It is well-known that along with the huge increase in neuronal RFs’ sizes within various visual cortices of the ventral visual pathway, from V1 to Inferior Temporal (IT) cortices, the complexity of encoded visual features increases hugely: from orientation selectivity in V1 to the emergence of faces and objects in IT (Rousselet et al., 2004; Kravitz et al., 2013). In parallel, the huge increases of RF sizes in the dorsal visual stream from V1 to MST, enable the encoding of direction selectivity at low speeds in V1 to the encoding of complex flow motion and high velocities in MST. However, as velocity is a measure of both motion direction and speed, the question remains as to how MT and MST achieve their capacity to encode high velocities when V1 DS neurons lose their direction selectivity beyond the V1 cutoff velocity (29°/s)?

### Cortical generation of direction selectivity

Occam’s razor, or the law of parsimony, suggests that the simplest solution to a problem is often the best one. A celebrated example of Occam’s razor in vision research is the de novo generation of orientation selectivity in V1 as suggested by Hubel and Wiesel, which was later supported experimentally in cat V1 (Hubel & Wiesel, 1965; Ferster, 1987; Reid & Alonso, 1995). Because V1 simple cells in both cats and monkeys share similar structured spatiotemporal RFs (Hubel & Wiesel, 1965), it is now well accepted, though not universally, that the generation of a simple cells’ orientation selectivity in macaque V1 uses similar mechanisms, i.e. by summing spike inputs primarily from LGN parvocellular neurons with spatially aligned RFs, to build elongated simple-cell RFs with ON-OFF subzones that are most responsive to elongated bars or edges. In parallel, DS simple cells in primate V1 layers 4Cα and 6 receive predominantly LGN magnocellular excitatory inputs, before projecting to layer 4B complex DS cells (Hawken et al., 1988; Angelucci & Sainsbury, 2006; Nassi & Callaway, 2009). This direction selectivity could increase from the input to the output layers of V1 through a cross-laminar refinement (Dai et al., 2025). Direction selectivity signifies sequential retinotopic activations aligned in space and time, i.e. spatiotemporal motion information. Linear and non-linear cortical integrations of convergent LGN excitatory inputs have been revealed experimentally to be the cortical basis for the generation of V1 orientation and direction responses in cats (Hubel & Wiesel, 1965; Ferster, 1987; Reid & Alonso, 1995). Thus, a simple spatiotemporal energy model can also elegantly reproduce DS responses in V1 (Simoncelli & Heeger, 1998; Bradley & Goyal, 2008; An et al., 2012; An et al., 2014). Analogously, excitatory monotonic synaptic projections of LGN to DS simple cells have been recently demonstrated in rabbit V1 (Su et al., 2025). Similar to cats, the cortical mechanisms underlying V1 direction selectivity in mice comprise, firstly, a linear and then a second non-linear integration of LGN inputs (Reid et al., 1987; DeAngelis et al., 1993; Jagadeesh et al., 1997; Lien & Scanziani, 2018; Rossi et al., 2020). The direction of the delayed asymmetric inhibition and the shift in excitatory responses have been revealed extracellularly to align with the directionality of primate V1 simple cells (M. S. Livingstone, 1998). Direction selectivity in MT and MST areas is conservatively believed to be inherited from V1 complex cells (Born & Bradley, 2005; Churchland et al., 2005; Priebe et al., 2006). Indeed, direct projections from V1 complex cells to MT DS neurons have been previously demonstrated by electrical stimulation, contributing to this popular hypothesis (Shipp & Zeki, 1989; J. A. Movshon & Newsome, 1996). However, intracortical inhibitory circuits have been found to contribute to direction selectivity not only within insects and rabbit retina (Borst & Helmstaedter, 2015; Rasmussen & Yonehara, 2020) but also in V1 of cats and primates (Sillito, 1975, 1977; Sato et al., 1995; N. C. Rust et al., 2005b). Similarly, intracortical inhibition exerts a major influence on the degree of direction selectivity in layer 2/3 of ferret V1 by suppressing responses to the null direction of motion (Wilson et al., 2018). Consistent with these findings, an inhibition mechanism is believed to contribute to both direction- and speed-selectivity tunings in macaque MT (Mikami et al., 1986a). More recently the direction selectivity of postsynaptic neurons in mouse V1 was found to be unrelated to the selectivity of the presynaptic neurons, but rather correlated with the spatial displacement between excitatory and inhibitory presynaptic ensembles (Rossi et al., 2020). This is consistent with an earlier study demonstrating direction selectivity emerges anew in mouse V1 (Lien & Scanziani, 2018). The evidence for de novo generation of directionality in mice V1 is also consistent with the initiation of DS responses within the retinal circuit motifs of many visual animals (Barlow & Levick, 1965; Vaney et al., 2012; J. S. Kim et al., 2014). Taken together these cross-species results support the case that direction selectivity could be generated afresh in MT, as spatiotemporal substructures are a built-in feature of DS cell’s large RFs. The alternative hypothesis of feature inheritance for MT direction selectivity at high speeds, perhaps most clearly stated in Churchland et al., 2005 p.1243 as “*… it is possible that high preferred speeds are present but rare in V1 and that this small minority of V1 neurons projects heavily to MT*”, requires that this minority of V1 DS cells are sufficient to generate the robust broadband tuning for high speeds in MT and MST. These sparse specialized cells would need to sample way beyond the visual space boundaries of their classical RFs, and MT would then need to multiplex across the preponderant low-speed and sparse high-speed afferent streams. To our knowledge there is no experimental data at present to support this idea. While not inconceivable, it seems unlikely that the direction selectivity of the majority of MT cells relies on the spiking inputs of a very small minority of V1 DS cells preferring speeds no higher than 30°/s. The spatial limits for DS cells in V1 seems matches those in area MT (Churchland et al., 2005). This is because that the spatial limits were measured at 32°/s, which were around the cutoff velocity of V1, and the optimal velocity of MT DS cells. In fact, this is in contrast to the measurements where the spatial limits of MT DS cells were three times larger than those of V1 when both cutoff velocities were used for the two areas, respectively (Mikami et al., 1986b). These class of arguments have been previously dismissed for pattern motion in MT (J. A. Movshon & Newsome, 1996; Majaj et al., 2007), with Majaj et al., 2007 p.369 stating that “*… it does not seem reasonable to suggest that the small numbers of pattern cells in earlier areas directly create the much higher prevalence of pattern selectivity of MT*”. If it were the case that this small group were responsible, we may expect the response latency of slow-tuned MT DS cells would be shorter. However, our latency analysis revealed that there is no statistical difference for MT DS cells with their preferred speeds below and above 30°/s (85.63ms versus 86.06ms; Kruskal-Wallis test, p=0.99; N=57 and 118, respectively), suggesting the processing of fast velocity in terms of response latency is in the same range of those preferring lower motion speed.

In previous studies of directionality in V1 and MT the motion speeds used are generally optimal for V1 (Maunsell & Van Essen, 1983; Orban et al., 1986; Rodman & Albright, 1987; Lagae et al., 1993; Priebe et al., 2003; Priebe et al., 2006). The cutoff velocity based on DSI values has not been specifically examined and reported for V1, MT and MST along the motion hierarchy. This may have previously been missed because although V1 and MT are very different in their optimal velocities, the overall velocity tuning curves of V1 and MT are similar and could have also contributed to the assumption that the direction selectivity in MT is entirely inherited from V1. In this study, we not only mapped the velocity tuning curves for LGN, V1, MT and MST, but also how the DSI tuning curves changed along the hierarchy. Regardless of cell types (simple and complex, component, pattern and unclassified), the cutoff velocity of V1 is far below that of MT and MST, and so the direction selectivity in MT and MST at faster speeds must be generated separately from the V1 population. The question is not whether, but how?

### Subcortical generation of direction selectivity

Given the ethological importance of rapid motion detection for survival, direction-selective neurons emerge at the earliest stages of the visual system (Baden et al., 2020; Summers et al., 2021). In non-primates, direction selectivity originates from starburst amacrine cells (ACs), followed by four ON-OFF Direction-selective ganglion cell (DSGC) types in rodent and rabbit retina (Fried et al., 2002; Briggman et al., 2011; Dhande et al., 2015; Wei, 2018; Rasmussen & Yonehara, 2020). Besides optical flow, visuo-motor behaviors such as balance and posture also involve these circuits (Sabbah et al., 2017; Rasmussen & Yonehara, 2020). In primates, DS ACs and one ON-OFF DSGC subtype (recursive bistratified) have recently been identified in vitro, comprising ∼1.5% of ganglion cells (Y. J. Kim et al., 2022). Thus, direction selectivity arises at the first retinal synapses across species, underscoring its conserved role, though strategies differ with ecological niche. A central question remains how DSGCs connect to diencephalic and midbrain structures such as the LGN and the Superior Colliculus (SC).

In mice, DSGCs project to superficial dLGN where DS responses are sharpened (Marshel et al., 2012), and to superficial V1 via a dedicated circuit (Cruz-Martin et al., 2014). In rabbits, DS motion signals are reliably transmitted to cortex through selective thalamocortical connections (Su et al., 2025). These findings suggest two parallel retino-geniculo-cortical pathways in mice and rabbits: one carrying DS signals from DSGCs via dLGN to superficial V1, and another conveying non-DS information to deeper V1 layer 4 (Cruz-Martin et al., 2014; Dhande et al., 2015). The SC, a major visuomotor integration hub, is a dominant retinorecipient structure in mice, receiving ∼85% of retinal ganglion projections (Ellis et al., 2016; Shi et al., 2017) and mediating ethologically relevant behaviors (Goettker & Gegenfurtner, 2021). Notably, while DS responses originate in the retina, those in dLGN and SC are thought to be inherited from DSGCs (Shi et al., 2017; Rasmussen & Yonehara, 2020). However, DSGC projections and roles in primate SC remain largely unknown.

Primate LGN cells are generally non-DS (Wiesel & Hubel, 1966; Derrington & Lennie, 1984), though some New World monkeys show mild biases (Xu et al., 2002; Eiber et al., 2018), consistent with early macaque studies (Lee et al., 1979). Thus, retinal DS signals appear “lost” en route to LGN. However, a direct retinopulvinar-to-MT pathway has been identified in marmosets, essential for MT development after birth (Warner et al., 2012; Warner et al., 2015). This minority of DS retinal cells may thus seed early MT maturation in New World monkeys. Only ∼10% of LGN and collicular cells show any DS responses (Goldberg & Wurtz, 1972; Xu et al., 2002), likely derived from DSGCs as in non-primates. In Old World monkeys, LGN lacks DS tuning because its dominant midget and parasol inputs are non-DS. Equal responses to all motion directions, as shown in **Fig.3a** of our study, may benefit LGN outputs to V1 layer 4Cβ, which is neither orientation nor direction biased. Strong orientation and direction selectivity then emerge de novo in V1 of both primates and mice through asymmetric feedforward inputs and intracortical inhibition (Hubel & Wiesel, 1963; J. A. Movshon et al., 1978; Reid et al., 1991; Jagadeesh et al., 1997; M. S. Livingstone, 1998; Lien & Scanziani, 2018; Rossi et al., 2020). In sum, spatiotemporal integration underlies motion processing from retina to V1: DSGCs arise from AC inputs in the retina, and DS responses may be inherited or generated anew in V1 from DS and non-DS LGN inputs. Yet, how visual systems can integrate motion signals at velocities beyond subcortical and V1 cutoffs remains unexamined.

### A spatiotemporal energy model simulation

MT is the most studied cortical area for processing visual motion information in the dorsal visual pathway (Born & Bradley, 2005; Nassi & Callaway, 2009). It is well accepted that specific and asymmetric feedforward projections from LGN magnocellular ON and OFF channels for constructing spatially separated ON and OFF subzones within RFs of V1 layer 4Cα simple cells (Hubel & Wiesel, 1962; Reid & Alonso, 1995; M. S. Livingstone & Conway, 2003). The RF substructures and their 2-D spatial interaction maps of V1 DS cells have been measured (M. S. Livingstone & Conway, 2003). Similar substructures with V1 DS cell RF sizes or even finer are successfully mapped at various local locations within large RFs of MT DS cells (M. S. Livingstone et al., 2001). Physiologically, these spatiotemporal inseparable RF substructures with excitatory and inhibitory subzones, exhibiting slightly different temporal dynamics, are believed to be the foundation for the initiation of direction selectivity in macaque V1 (M. S. Livingstone, 1998; M. S. Livingstone et al., 2001; M. S. Livingstone & Conway, 2003). Simoncelli and Heeger developed a two-stage integration model (Simoncelli & Heeger, 1998), where each stage computes a weighted linear sum of feedforward inputs, followed by rectification and divisive normalization to simulate a wide range of physiological data from both V1 and MT. Mathematically, ten basic spatiotemporal filters or kernels were built up to replicate various RF substructures of V1 layer 4 simple cells (Freeman & Adelson, 1991; Simoncelli & Heeger, 1998). This two-stage integration model can accounts for a wide range of physiological data (Simoncelli & Heeger, 1998). By taking advantage of this model (Simoncelli & Heeger, 1998), we developed our cascaded spatiotemporal integration model with the addition of a model of the cortical area MST and with additional projections to MT from both the LGN and V1 cells (Sincich et al., 2004; Nassi & Callaway, 2009). These extra projections from both the LGN and from V1 cells to MT proved to be essential as eliminating either the LGN or the V1 cell inputs in the cascaded spatiotemporal integration model generated simulations that failed to match experimental data (**Supplementary Fig.8**). This is backed up by the fact that LGN cells respond vigorously at all velocities (**Fig.3a**) and V1 neurons retain significantly higher firing rates above their cutoff velocity when compared to spontaneous responses (**Fig.3e**). Regardless of whether an object is moving fast or slow, sequential retinotopic activations are initiated in the retina and are common features for all neuron types within all retinotopic organized cortices. This precise spatiotemporal retinotopic information is captured by our fully integrated model from LGN to MST along the dorsal visual pathway and is the fundamental basis for generating direction selectivity at each processing stage from V1 and beyond. We further confirmed that MT can generate its DS responses purely from V1 orientation selectivity (OS) cells, as this spatiotemporal retinotopic information is also available with V1 OS cells (**Supplementary Fig.9**). These results support the findings that within weeks of V1 lesions, many neurons in MT still respond in a direction selective way to oriented bars and gratings presented inside the scotoma (Rodman & Albright, 1989; Rosa et al., 2000; Azzopardi et al., 2003; Alexander & Cowey, 2009). Recent studies found that with perceptual training, human patients with V1 lesions can recover the ability to discriminate the direction of moving dot patterns (Huxlin et al., 2009; Das et al., 2014). Interestingly, the representation of motion coherence information in MT+ of striate-lesion patients is similar to the V1 responses of healthy controls (Ajina et al., 2015). This certainly suggests MT can and does compute velocity de novo even when its anatomically dominant inputs are removed.

It is important to note that our cascaded integration model is a feedforward excitatory model **(Fig.7ab)**. This cascaded feedforward model captures the spatiotemporal motion energy changes within RFs of DS cells at each processing stage across a wide range of velocities, irrespective of whether the preceding cortices are direction or non-direction selective (**Fig.8 and Supplementary Fig.7**). The feedforward excitatory model for building velocity units or filters is not only the focus of our current study, but also the foundation of many previous models investigating cortical physiology, computation and motion perception (Grossberg, 1988; S. Nowlan & Sejnowski, 1993; S. J. Nowlan & Sejnowski, 1995; Koechlin et al., 1999). Although the modeling component remains qualitative, these models highlight dynamic interactions within the populations of cortical neurons between feedforward and intracortical recurrent connections underlying various motion perceptual phenomena. For example, the selection model emphasizes the most relevant motion of objects by globally extracting feedforward motion signals (S. Nowlan & Sejnowski, 1993; S. J. Nowlan & Sejnowski, 1995), whereas the Bayesian inference model relies on lateral recurrent excitation and inhibition to compute accurate local motion signals (Koechlin et al., 1999). Both are circuit-level models that use velocity units or filters as the initial feedforward stage, but neither addresses how these units encode the full range of motion speeds. Population decoding approaches, first developed in the motor system (Georgopoulos et al., 1986), have since been successfully applied to MT and MST, helping to account for motion responses and related behaviors (Deneve et al., 1999; Lisberger & Movshon, 1999; Pouget et al., 2000; Jazayeri & Movshon, 2006; Gu et al., 2010; Quaia et al., 2022; Behling & Lisberger, 2023; Lange et al., 2023; Levi et al., 2023; Mathis et al., 2024). We propose that more extensive population models, ideally combined with dense simultaneous recordings in V1, MT, and MST, will be crucial for revealing how spatiotemporally patterned activity shapes the emergence of motion features. Although the precise rules governing afferent pooling at each stage remain unresolved, the rich tapestry of direction-selective mechanisms found in nature provides fertile ground for future investigation.

**Fig. 8.**
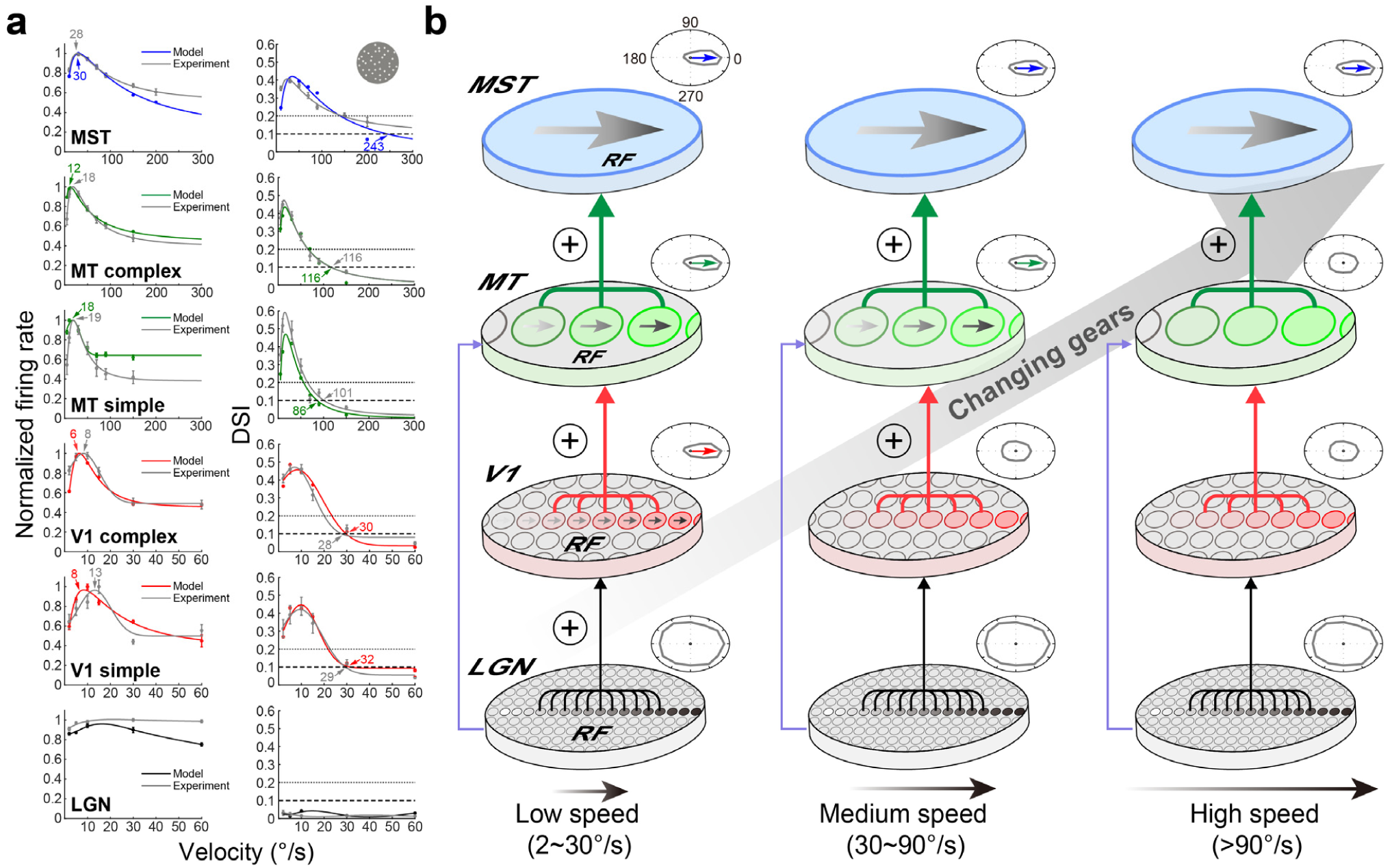
The computation and transformation of velocity across four successive cortices along the primate dorsal visual pathway. **a.** The velocity and DSI tuning responses across LGN, V1, MT and MST populations are reproduced by the cascaded spatiotemporal integration model. **b.** ‘Changing gears’, to automatically encode gradations of velocity along the hierarchy. The diagram summarizes hierarchical processing from the LGN to MST at varied speeds of motion; each stage integrates raw spikes initiated by sequential visuotopic activations from the preceding stages, with the progressive increase of RF sizes across cortices accommodating spatiotemporal integration at larger and larger scales, thus efficiently and automatically extracting direction of motion over the full range of perceived speeds. The plus symbol within the circle indicates linear and nonlinear integrations of feedforward spatiotemporal information. Direction tuning curves, oval or circular, respectively indicate the presence or absence of direction selectivity at each processing stage.

## Concluding remarks

Although we were not be able to make simultaneous multi-areal recordings in the dorsal visual stream, our findings by recordings from four successive visual areas, respectively, demonstrate that the primate motion pathway can accommodate a wide range of natural velocities through a *de novo* synthesis strategy that exploits the dramatic increase in RF size of DS neurons across the cortical hierarchy. Experimental results together with cascaded spatiotemporal integration model simulations spanning four successive dorsal stream areas, suggest that DS neurons can integrate sequential visuotopic inputs from earlier stages to generate velocity responses from LGN through MST. In essence, all DS neurons with spatiotemporally inseparable RFs function as efficient motion-energy detectors. At each stage, linear and non-linear integration of spatiotemporal inputs acts as an “automatic gear”, driving the progressive computation and transformation of velocity from low to high (**Fig.8b and Supplementary Fig.7b**). While motion may be processed by different organs in the vast array of nature’s sighted animals, the spatiotemporal information derived from sequential retinotopic/visuotopic activations generated by moving objects is essentially the same for all DS neurons to process. The law of parsimony suggests that the cortical mechanism that underlies the generation of direction selectivity at each computation stage in the primate dorsal visual pathway is not necessarily any more complex than that found in other lower species such as insects and rodents.

## METHODS

### Ethical approval

All primate experimental procedures were approved by the Animal Care and Use Committee of the Institute of Neuroscience, Chinese Academy of Sciences (No.ER-SIBS-221204P). All experimental procedures were also in accordance with the National Institutes of Health Guide for the Care and Use of Laboratory Animals.

### Neurophysiological experiments on nonhuman primates

#### Animal preparation

Four male rhesus monkeys (*Macaca mulatta*), aged 6 to 13 years and weighing 7∼12 kg were used in this study. A head post made of titanium alloy was initially implanted under sterile conditions for the head stabilization of each monkey. Monkeys were trained to perform passive fixation tasks for awake electrophysiology. The training procedure is identical to a previous study (Luo et al., 2019). We conducted the chamber installation operation only when the monkey had been well trained. By referring to their MRI scan, the plastic polyetheretherketone (PEEK) chamber was implanted over the position of LGN, V1, MT and MST (McAlonan et al., 2008; Kar & Krekelberg, 2016; Luo et al., 2019). The precise locations of LGN, MT and MST were determined based on an additional MRI scan with an electrode in situ (**Supplementary Fig.1a**).

#### Visual stimuli

Both moving random dots and drifting sine-wave gratings are classical motion stimuli. Velocity can be derived from SF/TF with drifting sine-wave gratings in macaque V1 and MT (Perrone & Thiele, 2001; Priebe et al., 2003; Priebe et al., 2006). The velocity can be directly measured with moving dots’ stimuli. We also employed natural movie stimuli to test MT and MST velocity responses.

Stimuli were generated with MATLAB (MathWorks) and Psychophysics toolbox 3 (Kleiner et al., 2007), displayed on a CRT monitor (Hewlett-Packard HSTND-1N01, 100 Hz). We used random dot stimuli (dot diameters and densities, 0.1° and 3.3 dot/°^2^ for LGN and V1, 0.3° and 1 dot/°^2^ for MT and MST, respectively). We also tested the smaller dots and bigger dots for MT and MST responses, there was no significant difference (p>0.32, N=33 for MT; p>0.65, N=38 for MST). In each trial, the dots moved in one of 12 equally stepped directions (0∼330°). We included a blank condition presented randomly to estimate spontaneous activity. The velocities (2∼60°/s, 5∼150°/s and 10∼200°/s) were used for LGN and V1, for MT and for MST, respectively. Each condition was randomly repeated at least 5 times.

Some DS neurons were also examined with drifting sine-wave gratings and plaid stimuli for the classification of simple and complex cells (Skottun et al., 1991), and component and pattern cells (J. Movshon et al., 1985; Khawaja et al., 2009), respectively. First, we identified the preferred direction of each cell by its strongest responses to gratings drifting in one direction of 12 equally stepped directions (0∼330°). Next, we tested a series of SF values (0.03125, 0.0625, 0.125, 0.25, 0.5, 1, 2, 4, 8 cycle/°) with a TF that yielded a robust response to determine the optimal SFs of DS neurons. Subsequently, we fixed the optimal SF and tested a range of TF values (0.1, 0.2, 0.4, 0.8, 1.6, 3.2, 6.4, 12.8, 25, 50 Hz) to determine the optimal TFs of DS neurons. We generated plaid stimuli by combining two gratings with optimal SF and TF, interleaved at 120°, moving in 12 different directions. We also used natural images of trees and leaves moving at various directions and velocities as additional stimuli for testing MT and MST. The direction and velocity settings were identical to those of random dots.

The total fixation time was 2000 ms for all stimuli above, with a 300 ms pre-stimulus time, a 1500 ms stimulus presenting time, and a 200 ms post-stimulus time. During the pre-stimulus and post-stimulus periods, no stimulus was presented and the monitor was set with a median gray background. The distance between the eyes of the monkey and the screen was 57 cm; the size of the screen was 1600×1200 pixels, subtending a visual angle of 30°×40°.

#### Electrophysiological neural recordings

The setups and procedures in awake monkey electrophysiological recording of MT and MST were previously reported^1^. In this study, we recorded LGN, V1, MT and MST with the same setups and procedures. Only neurons with their maximum direction-selective indexes (DSI) greater than 0.1 were recorded in V1, MT and MST (**Fig.2**), regardless of recording layers (Dai et al., 2025). The RF of V1, MT, and MST was hand mapped using computer generated moving random-dot fields. For the LGN recording, we first performed the hand mapping procedure with flashed stimuli to roughly determine the RF location (restricting the area to 2°×2°). Subsequently, we used white noise to precisely determine the RF location and size. LGN, V1, and MT rather than MST units exhibit a strong linear positive relationship between retinal eccentricity and RF diameter, consistent with previous studies (**Fig.2**)(Van Essen et al., 1984; Tanaka et al., 1986; M. Livingstone & Hubel, 1988; Celebrini & Newsome, 1994).

The RF diameter of LGN, V1, MT, and MST is <1.5°, 0.8∼2.1°, 2∼20.5°, 12∼50°, respectively. These RF sizes are in line with previous reports (Tanaka et al., 1986; Lagae et al., 1994; Cavanaugh et al., 2002; Perge et al., 2005; Jones et al., 2015; Luo et al., 2019).

### Data analysis

All electrophysiological data analysis and statistical calculations were performed using MATLAB. Neuron responses to moving stimuli were quantified by calculating the average firing rate within a 1500 ms window following stimulus onset. Neuronal response curves were quantitatively analyzed using a vector summation algorithm (Leventhal et al., 1995; An et al., 2014), the formula for which is:

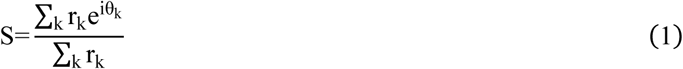

Here, θ_k_ represents the stimuli’ moving direction and r_k_ represents the average firing rate. S is a vector which angle represents the preferred direction, and the length represents the DSI. We only kept neurons of V1, MT and MST with clear direction-selective responses with DSI above 0.1 for further analysis (Schmolesky et al., 2000; Hua et al., 2006). Neurons with DSI≥0.1 at least at one tested velocity were considered to be DS neurons.

The average firing rate of the preferred direction and DSI at each velocity were fitted using the log-Gaussian function via the least squares method except for LGN (used smoothing: spline because of non-tuning). The log-Gaussian function is:

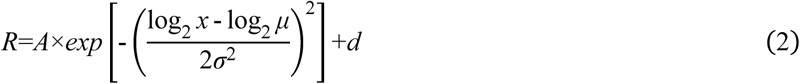

Where *R* represents the fitted firing rate or DSI, *A* represents the amplitude, *μ* represents the optimal velocity or optimal DSI velocity, *σ* represents the standard deviation, and *d* represents the offset. The velocity at which the firing rate is the highest was defined as the optimal velocity, and the velocity at which DSI decreases to 0.1 was defined as the cutoff velocity. The optimal velocity of some neurons was below the lowest velocity or above the highest velocity used in the experiment, in these cases the optimal velocity of those neurons was set to be the lowest velocity or the highest velocity in the experiment, respectively. The DSI of some neurons at 300°/s remains above 0.1, in these cases the cutoff velocity of those neurons was set to be the 300°/s maximum. The bandwidth is full width at 3 quarters maximum of the curve (the length of the range of velocities in which R≥0.75A+d). The population velocity tuning curve is the averaged response that normalized by the maximum of all neurons in each cortical area, with the baseline subtracted except for the model comparison in **Fig.8**. There is no statistical difference for optimal or cutoff velocity between the moving dots field and natural movie in MT and MST (p>0.5, Kruskal-Wallis test, N=512 and 39 for moving dots and natural stimuli, respectively).

We used the log-Gaussian function to fit the optimal SF and optimal TF similar to fitting the optimal velocity of random dots. V1 simple and complex cells can be best described with more linear filters (Nicole C. Rust et al., 2005a), and MT cells with a cascade linear-nonlinear integration model (Nicole C. Rust et al., 2006). Therefore, based on the linearity in response to drifting sine-wave gratings, we classified simple and complex cells in V1, MT and MST along the visual pathway. We computed the response linearity at different SF by calculating the response to drifting gratings. By obtaining the peri-stimulus time histogram (PSTH) within the 0.2∼1.5 s stimulus duration using a 25 ms time window (excluding the initial 0.2 s to remove the onset response), we used Fourier analysis to measure the first harmonic amplitude F1 and mean firing rate F0 from the PSTH for each SF (Y. Chen et al., 2009). Then we chose the max F1/F0 of the 3 most preferred SFs. The cell was classified as a simple cell if it was above 1, otherwise, the cell was classified as a complex cell. Among our populations across three cortices only in response to drifting sine-wave gratings, we noticed a small fraction of cells (about 15%) retains high responses along the increases of TFs at each cortex. These cells were mainly complex cells and included in all the analysis for the grating stimuli.

Partial correlation analysis was performed between neuron responses to moving gratings and plaids stimuli and the predictions of the pattern direction-selective (PDS) and component direction-selective (CDS) following previous methods (J. Movshon et al., 1985; Smith et al., 2005). The formula for calculating the partial correlation coefficient is:

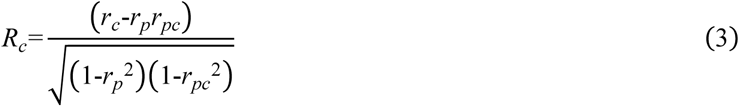

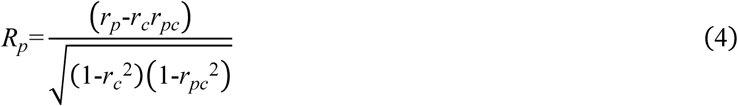

Where *R_c_* and *R_p_* represent the partial correlation coefficients between the neuron’s direction tuning curve and the predictions of the CDS and PDS models, respectively; *r_c_* and *r_p_* represent the correlation coefficients between the neuron’s direction tuning curve and the predictions of the CDS and PDS models, respectively; and *r_pc_* represents the correlation coefficient between the two prediction curves. Since the r-values did not follow the Gaussian distribution, the transformation from r-value to Z-score was performed for variance stabilization (Smith et al., 2005), using the following formula:

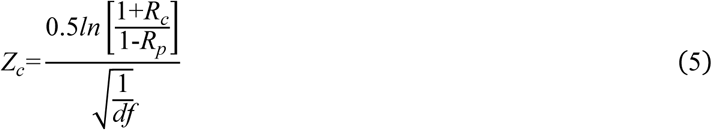

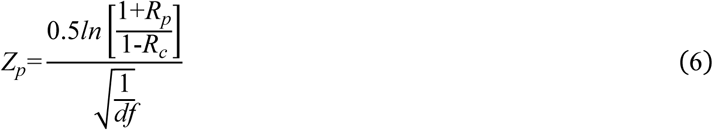

Where *df* is the degree of freedom. Cells were classified as pattern direction-selectivity (PDS) cells if *Z_p_*- *Z_c_*≥1.28. Similarly, cells were classified as component direction-selectivity (CDS) cells if *Z_c_*-*Z_p_*≥1.28. The remains were classified as unclassified cells.

We used white noise to determine the receptive field of the LGN. In the 1000 ms trial, the white square would randomly appear in the 10×10 matrix. We drew a heat map of the cell’s response to each square position. After applying two-dimensional Gaussian smoothing to the heat map, we obtained the receptive field’s position and diameter.

Parvocellular cells (P cell) and magnocellular cells (M cell) of LGN show different responses to spots of a flash of white light or colored light. Most P cells were clearly color opponent, while M cells had broad-band characteristics or showed some slight selectivity for wavelength without color opponency (Schiller & Malpeli, 1978). We used flashes of different colors with color opponency to distinguish P cells and M cells. There were 20 (34.48%) M cells and 38 (65.52%) P cells of the LGN cells we recorded.

Differences among groups were analyzed using the Kruskal-Wallis test. If the test indicated significant differences and more than two groups were tested, multiple comparison correction was performed using the Bonferroni method. Linear relationships were assessed using Spearman rank correlation. All tests were analyzed using MATLAB. All statistical results were considered significant at a level of p < 0.05. All error bars were SEM.

### Hierarchical velocity computation model

#### Cascaded spatial-temporal model for dorsal visual pathway

Several computational models were previously constructed to study motion coding in MT. Nowlan and Sejnowski 1995’s model integrates velocity inputs directly from V1 (S. J. Nowlan & Sejnowski, 1995), whereas the model by Simoncelli and Heeger 1998 takes neural responses of V1 as inputs to compute the velocity (Simoncelli & Heeger, 1998). Both models did not consider the response properties in V1 and MT at high velocity and the contribution of LGN direct inputs to MT. In our study, the model was constructed with multiple stages/components, corresponding to hierarchical visual areas of LGN, V1, MT and MST in the visual dorsal pathway (**Fig.1b**). Each cortical stage (V1, MT, and MST) receives neural response outputs from its previous stage through spatial-temporal filters. The filters were adopted directly from the SH model (Simoncelli & Heeger, 1998). We set the fitting constraints for the receptive field sizes based on our experimental results (V1: ∼2°, MT: ∼7°, MST: ∼29°). In the model, the component for V1 was driven by pre-cortical output (from LGN); the component for MT was driven by both V1 and pre-cortical outputs (from LGN); and the component for MST was only driven by MT output. The connection weights between different brain regions and the ratio of simple to complex cells in V1 were initially set according to the existing literatures (Freeman & Adelson, 1991; Simoncelli & Heeger, 1998; Sincich et al., 2004; Bradley & Goyal, 2008; Nassi & Callaway, 2009). These connection weights were then adjusted to best fit the experiment data by varying the relative connection strengths for LGN to MT and V1 to MT, respectively. The details of the model components are described as follows.

#### Visual stimuli (drifting gratings and moving dots)

The visual stimuli, *S*(*x*,*y*,*t*), for model simulations include drifting sine-wave gratings and moving dots’ texture. Both types of stimuli are created in a three-dimensional matrix *S*(*x*,*y*,*t*) (in equation 7) which contains luminance values (0∼1) at different locations (*x*,*y*) and time points *t*. The spatial locations have 102 pixels for V1, 357 for MT and 1182 for MST. The time is in the temporal resolution of 10 ms (corresponding to one frame duration of a screen with frame rate at 100 fps) and lasts for more than a full temporal cycle (4 seconds for TF=0.25 Hz) in drifting grating and 1.5 seconds (150 time points) in moving dots.

#### The stage for pre-cortical processing and thalamic-cortical transmission (thalamic drive)

In the model, we used linear spatial-temporal filters, to mimic the pre-cortical processes (including dynamic response properties of LGN) and thalamic-cortical transmission (Simoncelli & Heeger, 1998), which are defined as thalamic drives (equation 7) to V1 and MT. The convolutions of these spatial-temporal filters and visual stimuli (moving dots or drifting gratings) are sent to stages for V1 and MT with different spatial scales for filters (*H_V1_*(x,y,t) for V1 and *H_MT_*_1_(*x*,*y*,*t*) for MT) correspondingly.

The spatial-temporal filters, *H_c_*(*x*,*y*,*t*), are generated by a weighted sum of 10 basic spatiotemporal filters, *b_i_*(*x*,*y*,*t*), which are spatial-temporal separable filters (Freeman & Adelson, 1991; Simoncelli & Heeger, 1998).

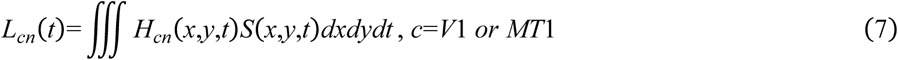

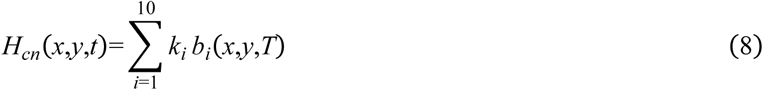

The LGN directly uses 10 basic spatiotemporal filters that have been processed by Rectified Linear Unit (ReLU) in spatial frequency. Based on this, the response of 10 LGN cells can be obtained. The response of the i-th LGN is

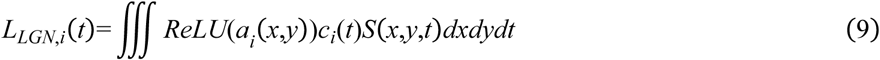

where *ReLU*(*a_i_*(*x*,*y*)) and *c_i_*(*t*) is the spatial filter and temporal filter of the i-th LGN.

#### Model V1

V1 stage in the model includes both simple and complex cells (Hubel & Wiesel, 1962). There are 12 simple neurons and 12 complex neurons with different direction preferences (evenly and circularly spacing between 0 and 360 degree), at V1 stage. Responses of each simple cell are generated by half-squared rectification (equation 10) of summed thalamic drives (*L_c_*) to this neuron followed by a normalization (equation 11). Responses of each complex cell are generated by full-squared rectification and blur (equation 12) of summed thalamic drives to this neuron followed by a normalization (equation 13). The computation of complex cells is equivalent to the summation of a pair of V1 simple cells with opposite phase preferences followed by normalization (Simoncelli & Heeger, 1998).

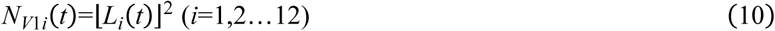

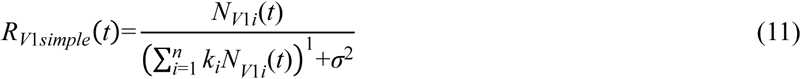

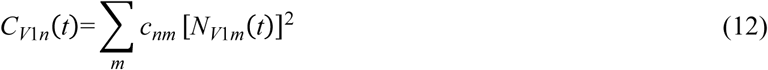

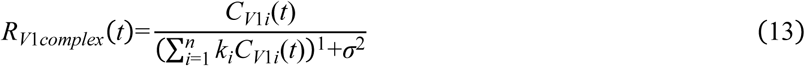

#### Model MT

MT neurons in the model are not only driven by outputs of V1 simple and complex cells, but also receives thalamic drives (equation 7). At MT stage, there are also 12 simple neurons and 12 complex neurons with different direction preferences (evenly and circularly spacing between 0 and 360 degree). For V1 drives to MT, both MT simple and complex cell responses go through spatial-temporal filters, *H_MT_*_2_(*x*,*y*,*t*) and *H_MT_*_3_(*x*,*y*,*t*) (equations 14 and 15); then the weighted (*w*_1_, *w*_2_ and *w*_3_ in equation 16) summation of the two V1 drives (*L_MT_*_2_ and *L_MT_*_3_) and thalamic drives (*L_MT_*_1_) are half-wave rectified (equation 16) followed by normalization (equation 17) for MT simple cell.

Responses of each MT complex cell are generated by full-squared rectification and blur (equation 18) and normalization (equation 19).

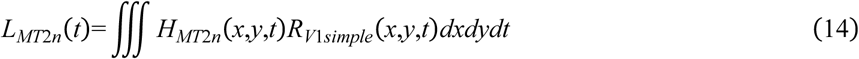

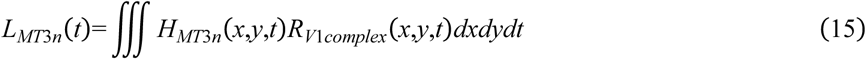

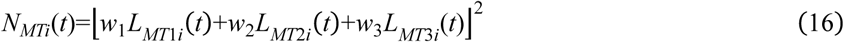

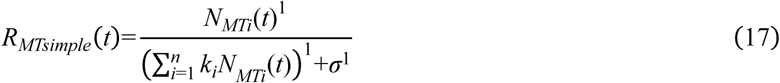

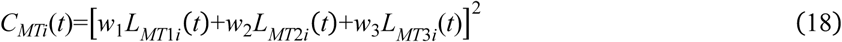

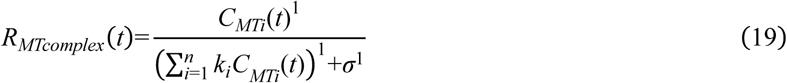

Model MT estimated contributions from inputs of LGN, V1 simple and complex cells at 25%, 8% and 67%, respectively for our simulated results.

#### Model MST

MST neurons in the model are only driven by outputs of MT simple and complex cells (equation 20). The MT drives to MST are computed by convolution of spatial-temporal filters (*H_MST_*(*x*,*y*,*t*)) and MT outputs followed by full-wave rectification and normalization (equations 22&23). The contributions of MT simple and complex cells to MST (*w*_4_ and *w*_5_ in equation 21) are 17% and 83% in our MST (*w*_4_ and *w*_5_ in equation 21) are 17% and 83% in our MST simulation, respectively.

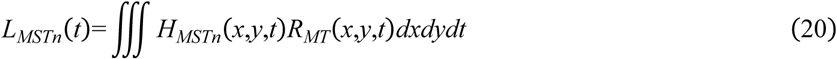

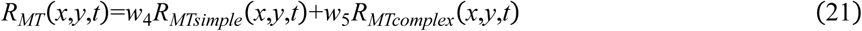

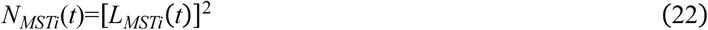

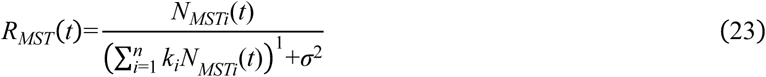

## Author contribution

K.Y.H., L.X.L., and J.X.L. performed experiments and data analysis, and Y.L.L., J.P.Y., Y.F.L., W.H.X., Y.Z.J.L., X.H.L., I.M.A., S.S., H.B.Y., N.M., and W.W. contributed to the data analysis. Y.W. and D.J.X. performed the model simulation. W.W., N.M. and I.M.A. wrote the main paper. W.W. designed and supervised the research.

## Supporting information

Graphical abstract & Supplementary figures

## Acknowledgement

This work was supported by the following grants: STI2030-Major Projects 2022ZD0204600 (to W.W. and D.J.X), the CAS Project for Young Scientists in Basic Research (Grant No. YSBR-113) (to Y.L.L), National Natural Science Foundation of China grants 32070992, 32150410370 (to I.M.A.).

## Supplemental information list

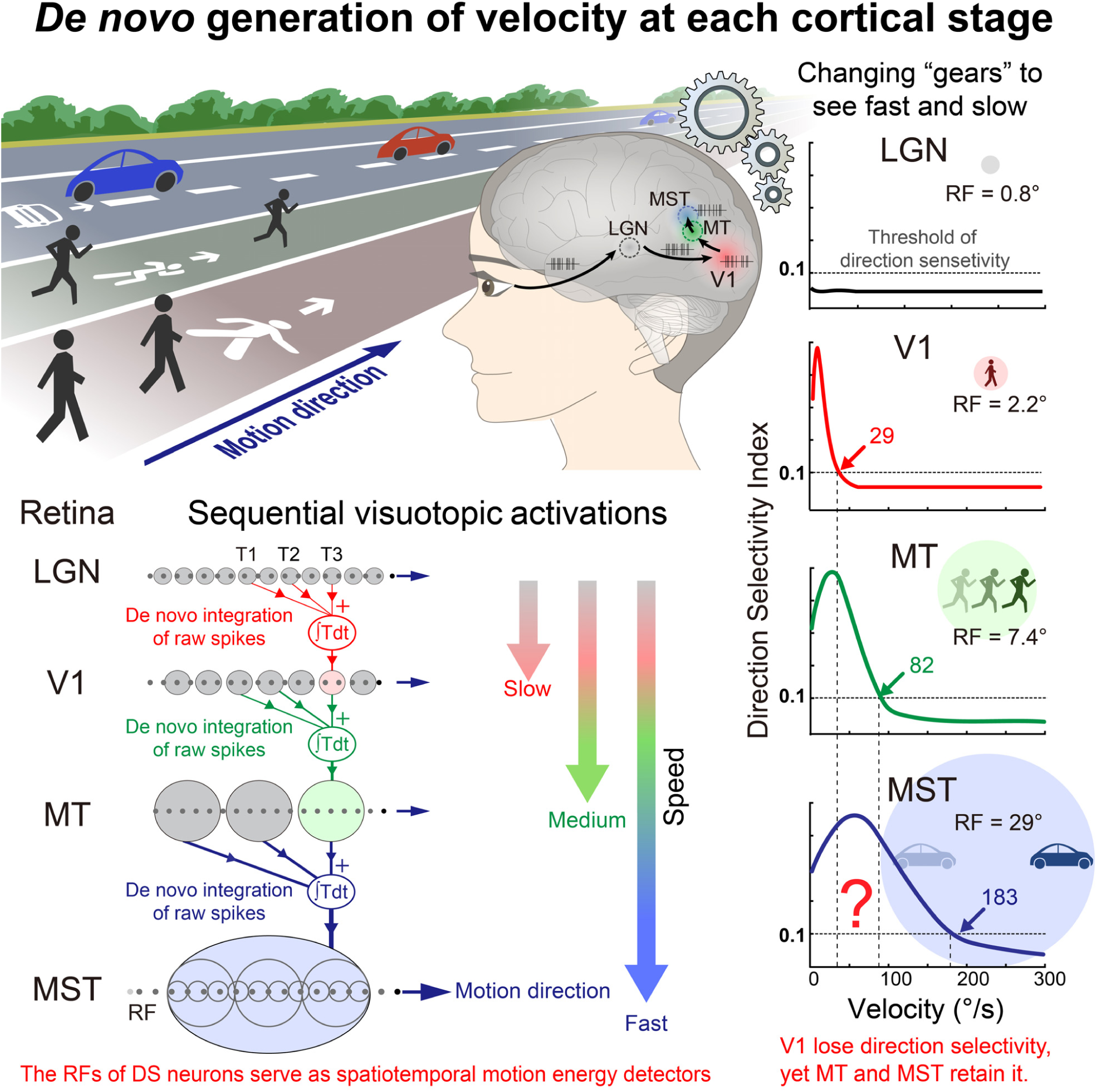

Graphic abstract shows how our hierarchically organized visual system employs each cortical area as a distinct "gear" to compute velocity anew for a wide spectrum of object speeds. The optimal and cutoff velocities of the tuning curves in V1, MT and MST are averaged velocities of recorded neurons across the three areas in awake macaques.

## Supplementary figures

**Supplementary Fig. 1.**
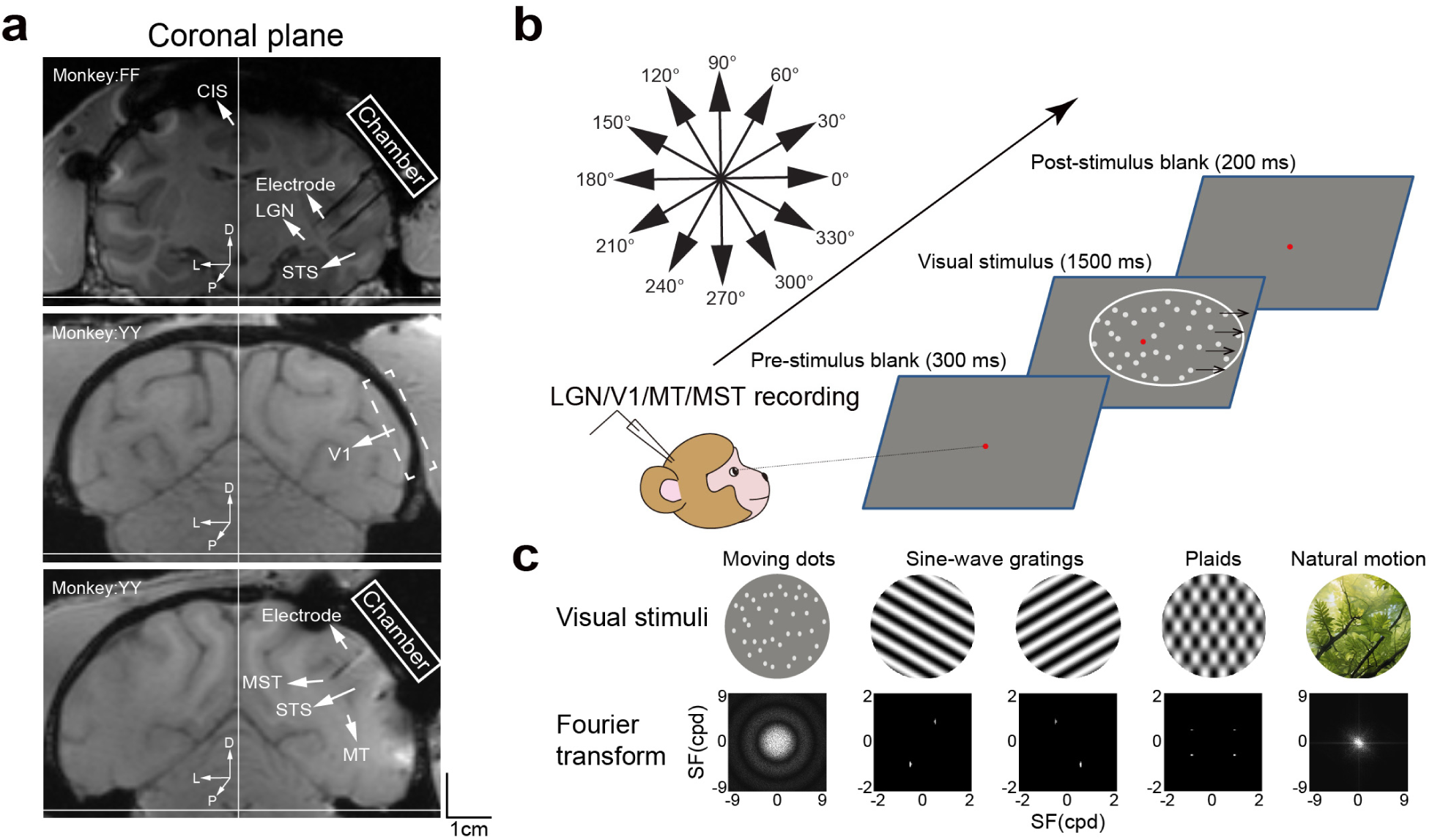
The experiment design, recording methods and visual stimuli. **a.** MRI localization of visual brain areas of the lateral geniculate nucleus (LGN), the primary visual cortex (V1), the middle temporal area (MT) and the medial superior temporal area (MST) preparing for electrode penetrations. CIS, cingulate sulcus; STS, superior temporal sulcus. **b.** Stimulation settings and visual neuron recordings in LGN, V1, MT and MST along the hierarchy of the dorsal visual stream. **c.** The visual stimuli and their SF distributions after Fourier transformation.

**Supplementary Fig. 2.**
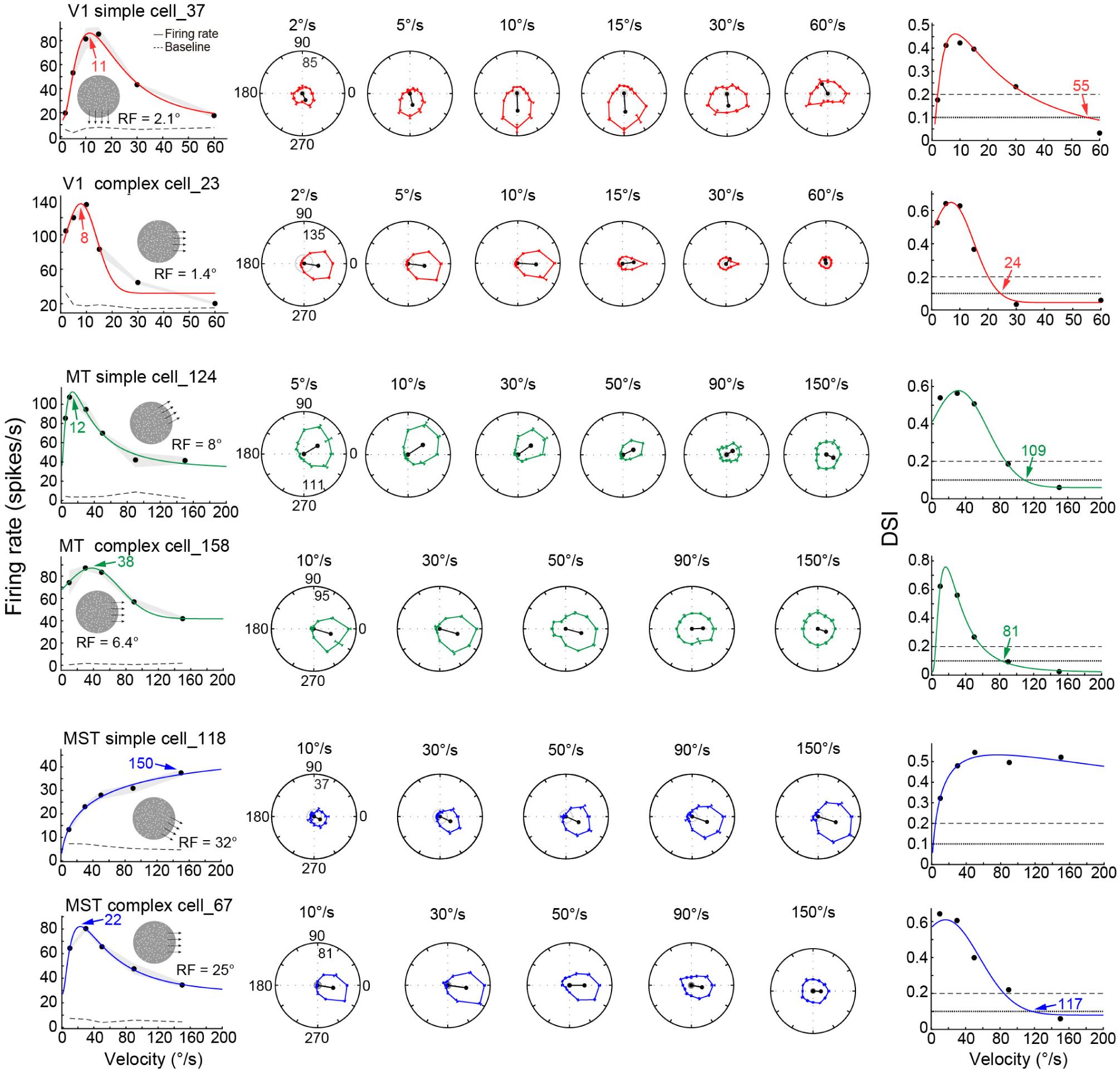
The velocity tuning examples of simple and complex cells in V1, MT and MST. The classification of these examples is presented in **Fig.4a**. The left column shows the velocity tunings of each cell type across cortices, while the middle column exhibits the corresponding direction circular tuning curves of each cell type. The shade and error bar indicate the SEM, and also for the rest figures. The right column illustrates the DSI tuning curves of the corresponding cell types in the left and middle columns. The colored digits with arrow points indicate the optimal and cutoff velocities, respectively, and also for the rest figures.

**Supplementary Fig. 3.**
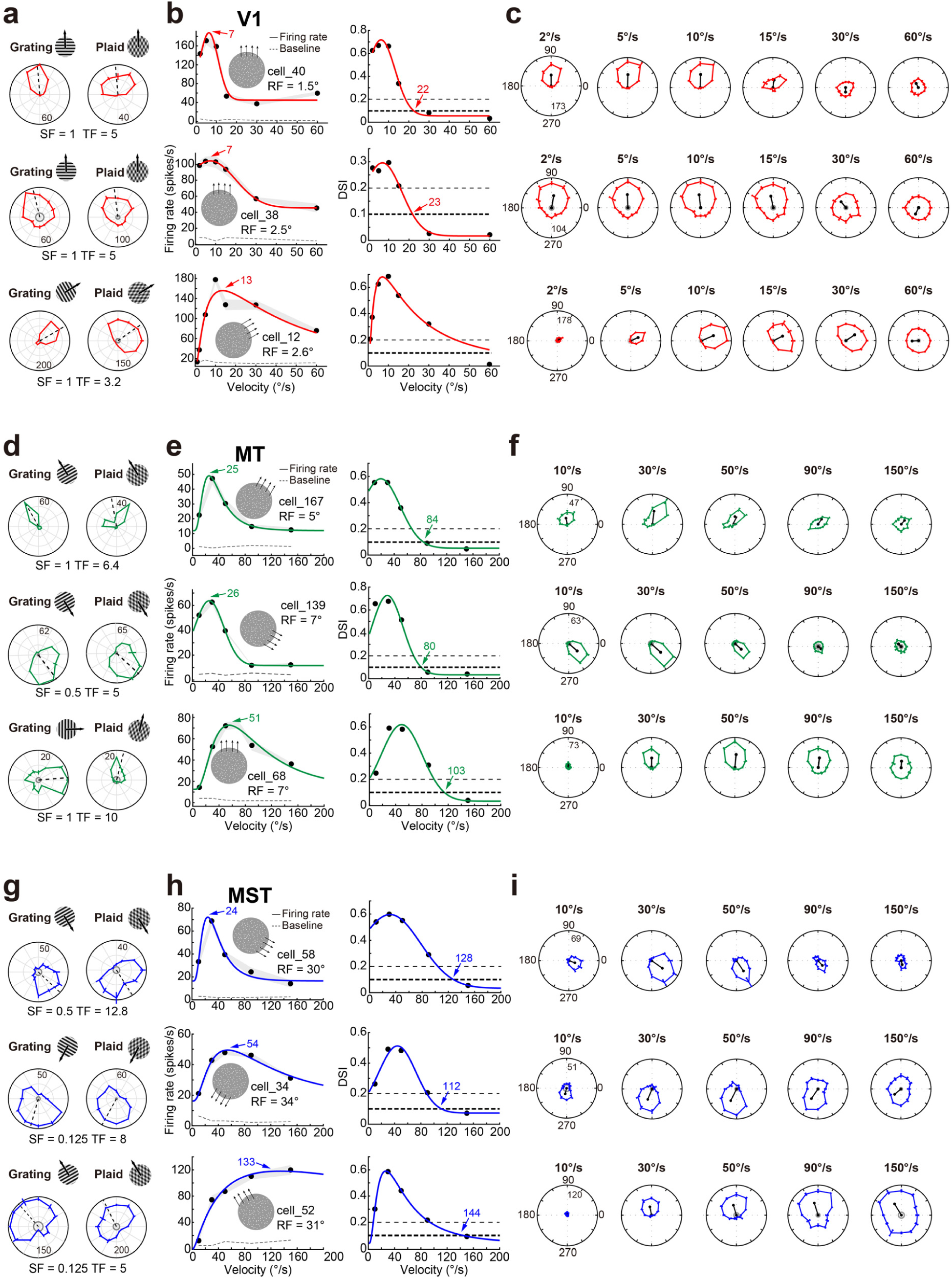
The component, pattern, and unclassified cell examples in V1, MT and MST. **a.** The classification of three cell types in V1. **b.** The velocity tuning responses and DSI tuning curves for examples of the three cell types. **c.** The corresponding direction tuning curves in circular plots. **d.** The classification of three cell types in MT. **e.** The velocity tuning responses and DSI tuning curves for examples of the three cell types. **f.** The corresponding direction tuning curves in circular plots. **g.** The classification of three cell types in MST. **h.** The velocity tuning responses and DSI tuning curves for examples of the three cell types. **i.** The corresponding direction tuning curves in circular plots.

**Supplementary Fig. 4.**
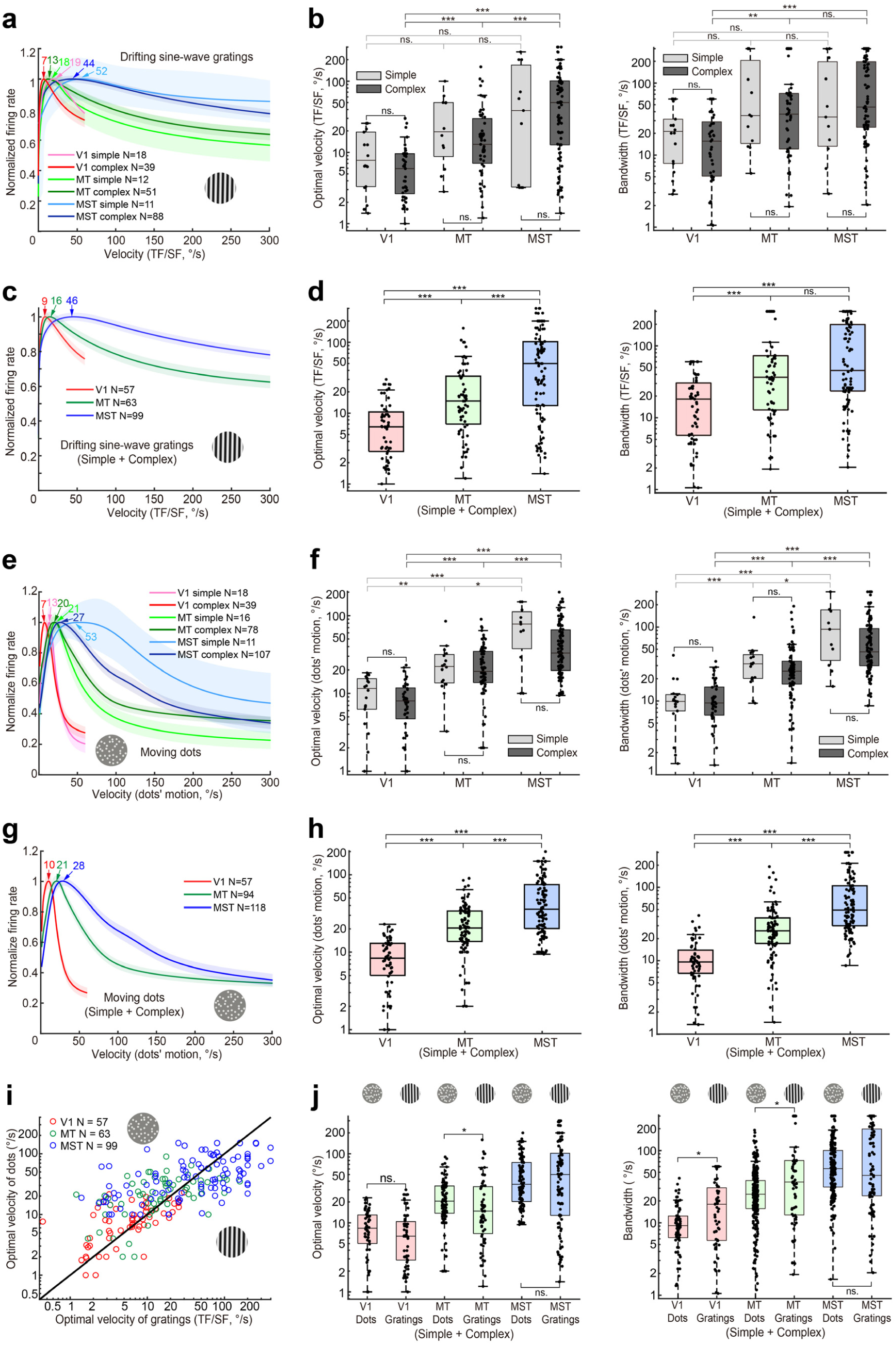
The velocity tuning responses of simple and complex cells to drifting sine-wave gratings and random dot motion across V1, MT and MST. **a.** The velocity tuning responses of simple and complex cells to drifting gratings across cortices along the hierarchy. **b.** The comparison of optimal velocities and bandwidths for simple and complex cells within and across cortices of V1, MT and MST. The optimal velocity of simple cells: 11.51±2.10°/s, 32.30±8.65°/s and 78.33±28.45°/s for V1, MT and MST, respectively. The optimal velocity of complex cells: 7.15±1.08°/s, 23.93±4.29°/s and 74.86±8.31°/s for V1, MT and MST, respectively. The bandwidth of simple cells: 25.20±4.85°/s, 75.23±13.71°/s and 98.24±35.74°/s for V1, MT and MST, respectively. The bandwidth of complex cells: 20.47±2.92°/s, 75.23±13.71°/s and 108.51±111.60°/s for V1, MT and MST, respectively. **c.** The velocity tuning responses without separating simple and complex cells. **d.** The comparison of optimal velocities and bandwidths of the populations across V1, MT and MST. **e.** The velocity tuning responses of simple and complex cells to moving dots. **f.** The comparison of the optimal velocity and bandwidth of simple and complex cells within and across V1, MT and MST. The optimal velocity of simple cells: 10.36±1.33°/s, 25.45±4.70°/s and 77.19±14.86°/s for V1, MT and MST, respectively. The optimal velocity of complex cells: 8.62±0.90°/s, 24.94±1.95°/s and 49.01±3.91°/s for V1, MT and MST, respectively. The bandwidth of simple cells: 11.31±2.13°/s, 34.44±4.57°/s and 120.29±31.17°/s for V1, MT and MST, respectively. The bandwidth of complex cells: 11.83±1.26°/s, 33.04±3.63°/s and 75.45±6.98°/s for V1, MT and MST, respectively. **g.** The velocity tuning responses without separating simple and complex cells along the hierarchy. **h.** The comparison of the optimal velocity and bandwidth of populations across V1, MT and MST. **i.** The comparison of optimal velocities between moving fields of dots motion and drifting sine-wave gratings within each cortex along the hierarchy. **j.** The comparison of optimal velocities and bandwidths between the motion of dots and sine-wave gratings within each cortex of V1, MT and MST, respectively.

**Supplementary Fig. 5.**
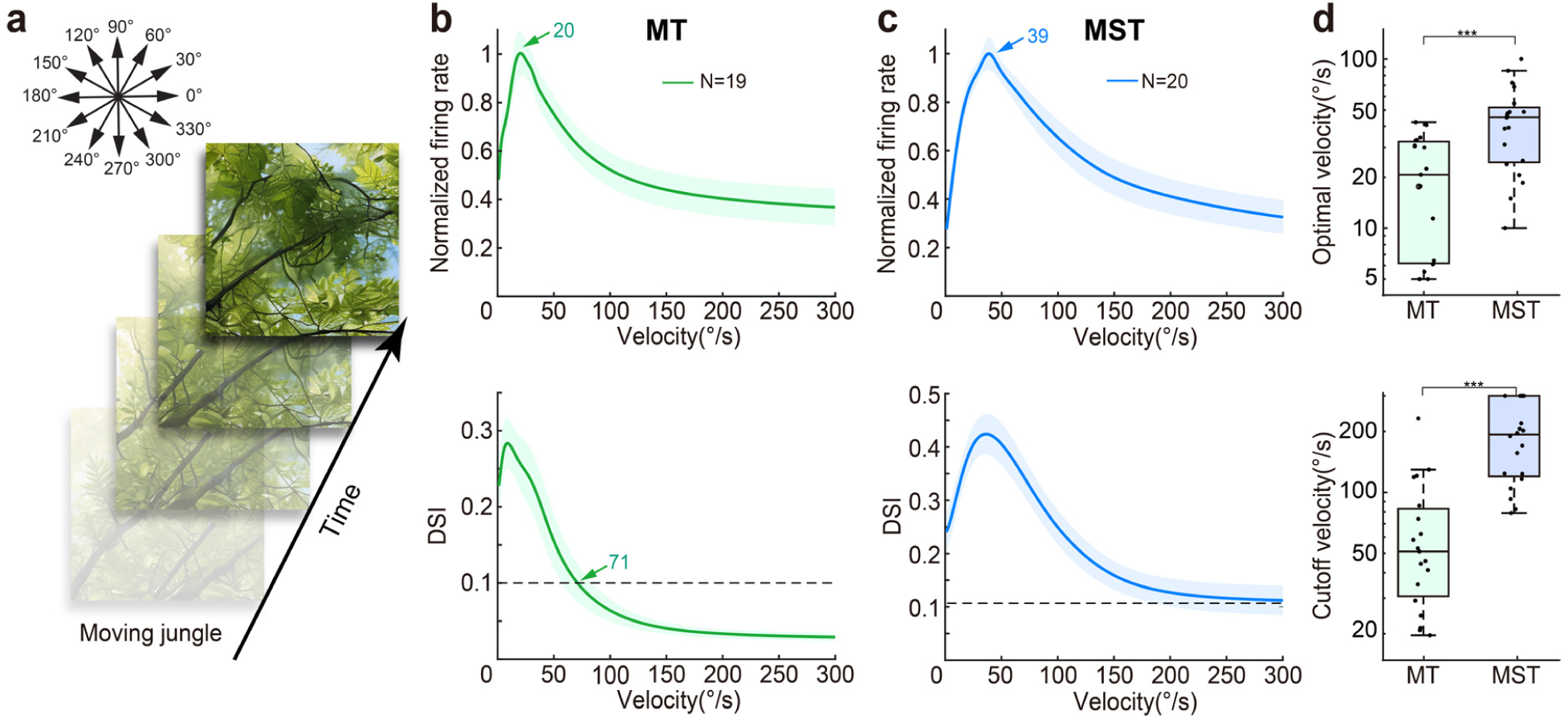
Natural stimuli for mapping MT and MST velocity and DSI tuning responses. **a.** Example frames from a natural movie of a jungle scene. The velocity of natural movies used ranged from 5 to 200 °/s, and the movies presented within RFs of both MT and MST. **b.** The velocity and DSI tuning curves of MT DS neurons. **c.** The velocity and DSI tuning curves of MST DS neurons. **d.** The comparisons of optimal and cutoff velocities between MT and MST. Similarly to dots motion, both the optimal and cutoff velocities in MST are significantly higher than those in MT by using stimuli of natural image movies. The optimal velocity: 20.96±3.03°/s versus 44.33±5.29°/s for MT and MST, respectively. The cutoff velocity: 66.72±12.10°/s versus 178.53±15.10°/s for MT and MST, respectively.

**Supplementary Fig. 6.**
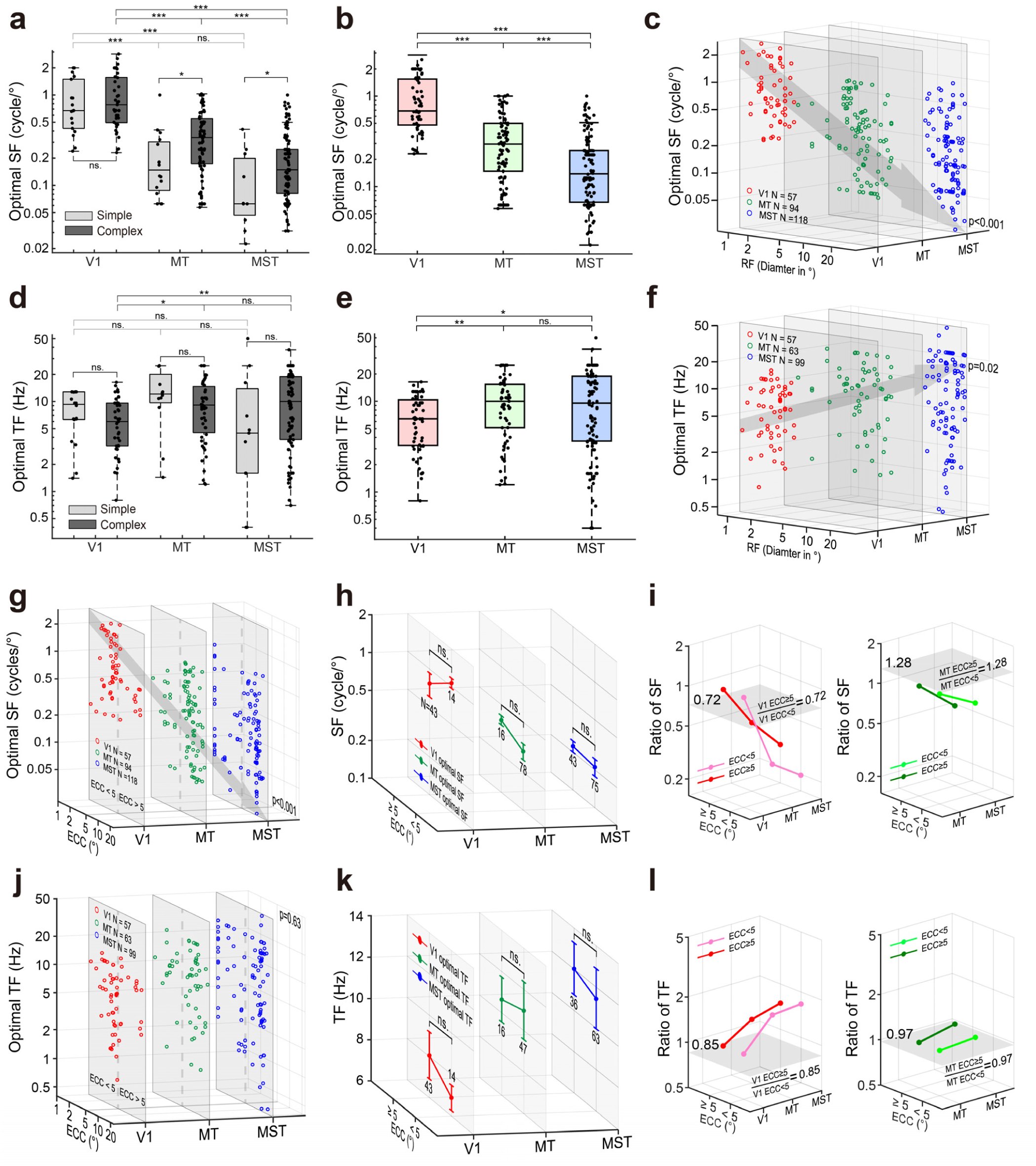
The comparison of optimal SFs and TFs cross cortices and eccentricities. **a.** The comparison of optimal SFs for simple and complex cells across V1, MT and MST. There is no significant difference for V1 complex and simple cells in their optimal SF, consistent with previous reports (Foster et al., 1985). **b.** The comparison of optimal SFs without separating simple and complex cells. The optimal SF:0.99±0.09 cycle/°, 0.36±0.03 cycle/° and 0.22±0.02 cycle/° for V1, MT, and MST, respectively. **c.** The optimal SFs are reversely correlated with the increases of RFs across V1, MT and MST. **d.** The comparison of optimal TFs for simple and complex cells across V1, MT and MST. **e.** The comparison of optimal TFs without separating simple and complex cells across V1, MT and MST. The optimal TF: 7.23±0.54 Hz, 10.73±0.87 Hz and 11.8±0.96 Hz for V1, MT, and MST, respectively. **f.** The optimal TFs of DS neurons are weakly correlated with the increase of RF diameters across V1, MT and MST. **g.** The optimal SFs are reversely correlated with the increases of eccentricities across V1, MT and MST. **h.** The comparison of optimal SFs between below and above 5° eccentricities, respectively, across V1, MT and MST. **i.** The relative changes of the optimal SFs across cortices after normalizing to V1 and MT, respectively. The gray surfaces indicate the ratio levels of the optimal SFs for neurons above and below 5° eccentricities for V1 and MT, respectively. **j.** No correlation between optimal TFs and eccentricity across V1, MT and MST. **k.** The comparison of optimal TFs between below and above 5° eccentricities, respectively, across V1, MT and MST. **l.** The relative changes of the optimal TFs across cortices after normalizing them to V1 and MT, respectively. The gray surfaces indicate the ratio levels of the optimal TFs for neurons above and below 5° eccentricities within V1 and MT, respectively.

**Supplementary Fig. 7.**
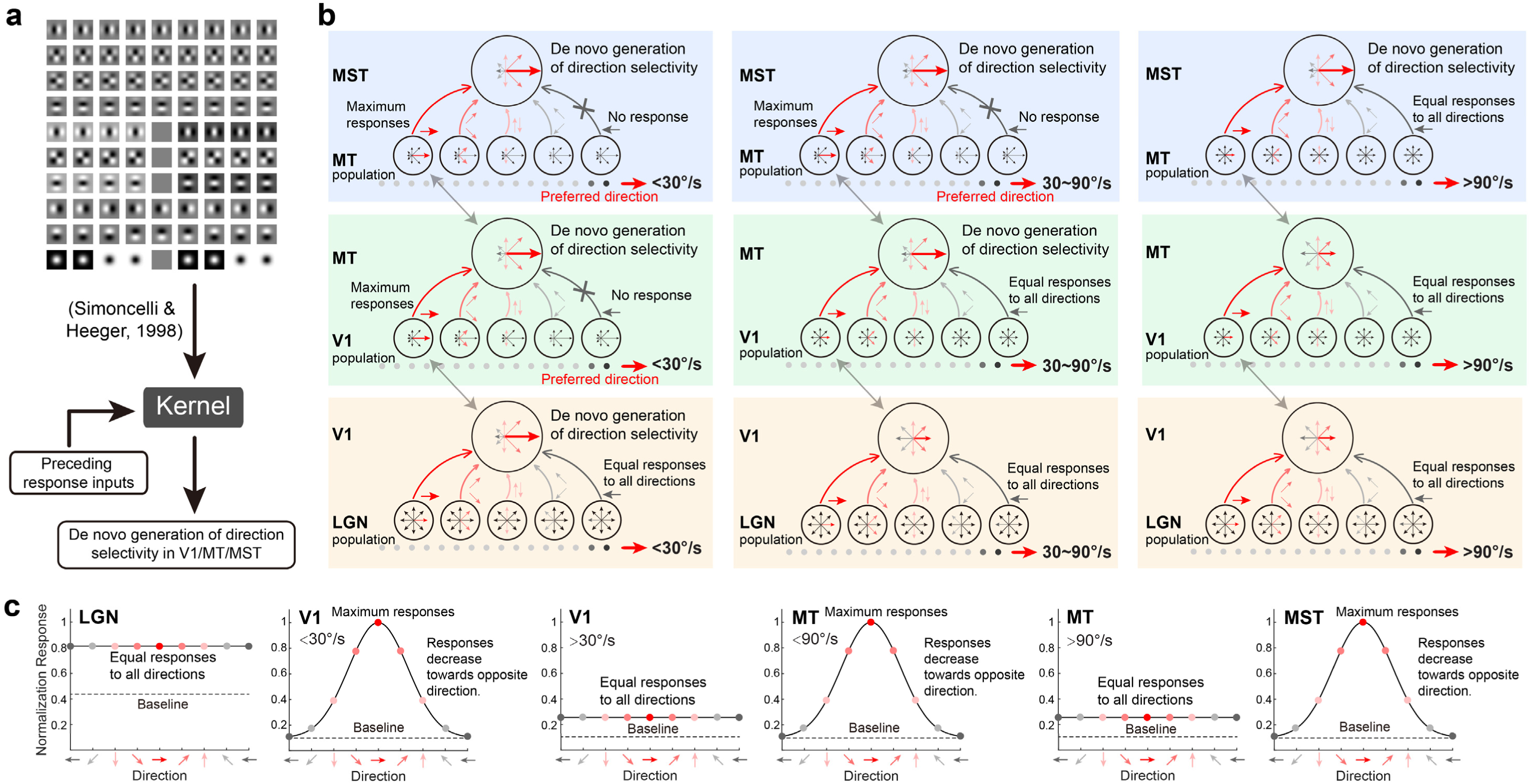
Schematic diagrams depict the cascaded spatiotemporal integration of presynaptic neuronal responses across cortices in the primate dorsal visual pathway. **a. The mathematic filters (or kernels) replicate spatiotemporal RF substructures of V1 DS cells.** Ten basic spatiotemporal filters have been built to replicate the RF structures of DS cells (Freeman & Adelson, 1991; Simoncelli & Heeger, 1998), which are the core foundation of our cascaded spatiotemporal integration model. **b. The direction selective responses across V1, MT and MST associated with motion speeds.** For given moving stimuli, presynaptic spikes are maximum only when the stimulus is with the DS neuron’s preferred motion direction. When the stimulus is away from the preferred direction, the firing rate decreases significantly. Direction selectivity varies across V1 and MT, respectively, which depending on different ranges of motion speeds. **c. The direction selectivity depending on the motion speed, in V1, MT and MST, respectively.** LGN cells respond equally well to all motion directions, while V1 and MT alternatively lost their direction selectivity at their cutoff velocities of above 30 and 90°/s, respectively (Fig.3, please also see Mikami et al., 1986b; Churchland et al., 2005). The cutoff velocity of MST DS cells is beyond what we can possibly measure due to the fresh rate limitation of the stimulus monitor. Both the normalized responses and width of the tuning curves are just for illustration only.

**Supplementary Fig. 8.**
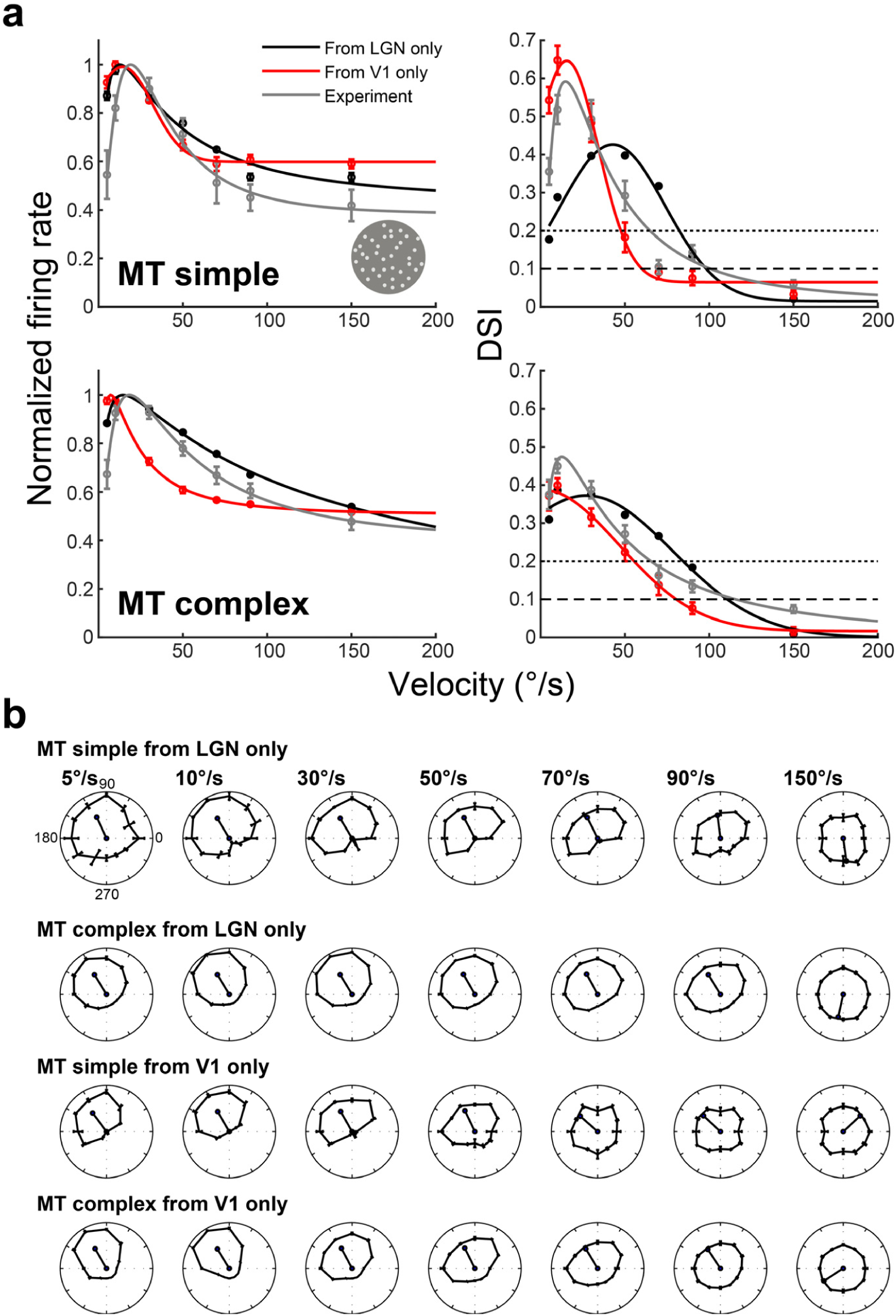
The model predicted results of velocity and DSI tuning responses in MT with and without V1 and LGN inputs, respectively. **a.** The CSTI model simulated velocity and DSI tuning responses of MT simple and complex cells after blocking V1 and LGN inputs, respectively. **b.** The CSTI model simulated direction circular tuning curves at each speed for simple and complex cells in MT, after blocking LGN and V1 inputs, respectively.

**Supplementary Fig. 9.**
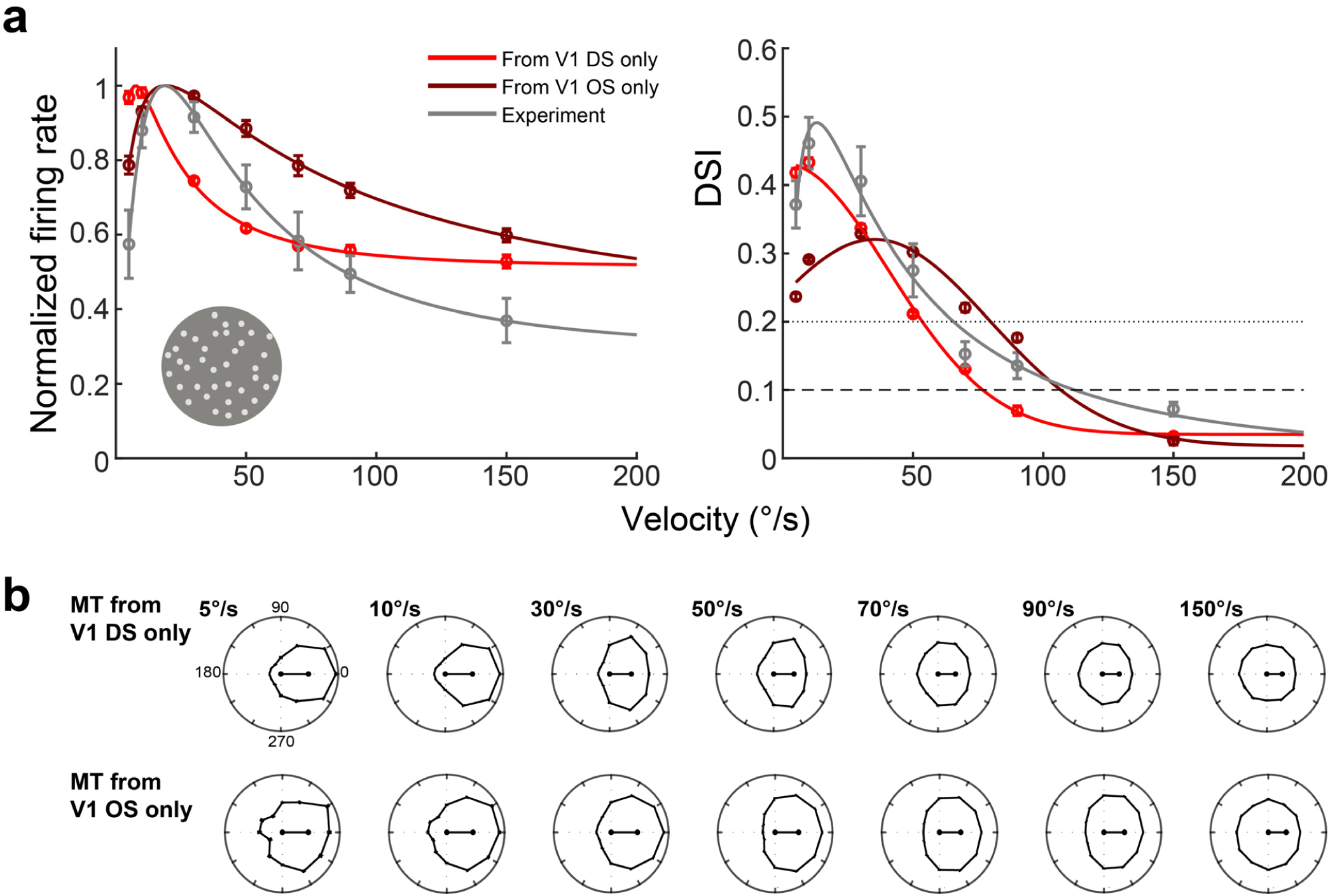
The model predicted results of velocity and DSI tuning responses in MT with only V1 DS and V1 OS inputs, respectively. **a.** The CSTI model simulated velocity and DSI tuning responses of MT with V1 DS and V1 OS inputs only, respectively. **b.** The CSTI model simulated direction circular tuning curves at each speed for MT, from V1 DS and V1 OS inputs only, respectively. MT can generate its DS responses de novo irrespective of its presynaptic inputs.

