## Supplementary material for "Beyond Inheritance: De novo Fast Motion Computation in Primate Visual Cortex": Graphical abstract & Supplementary figures

**Supplemental information list:**

1. **Graphical abstract**
2. **Supplementary figures 1~9**

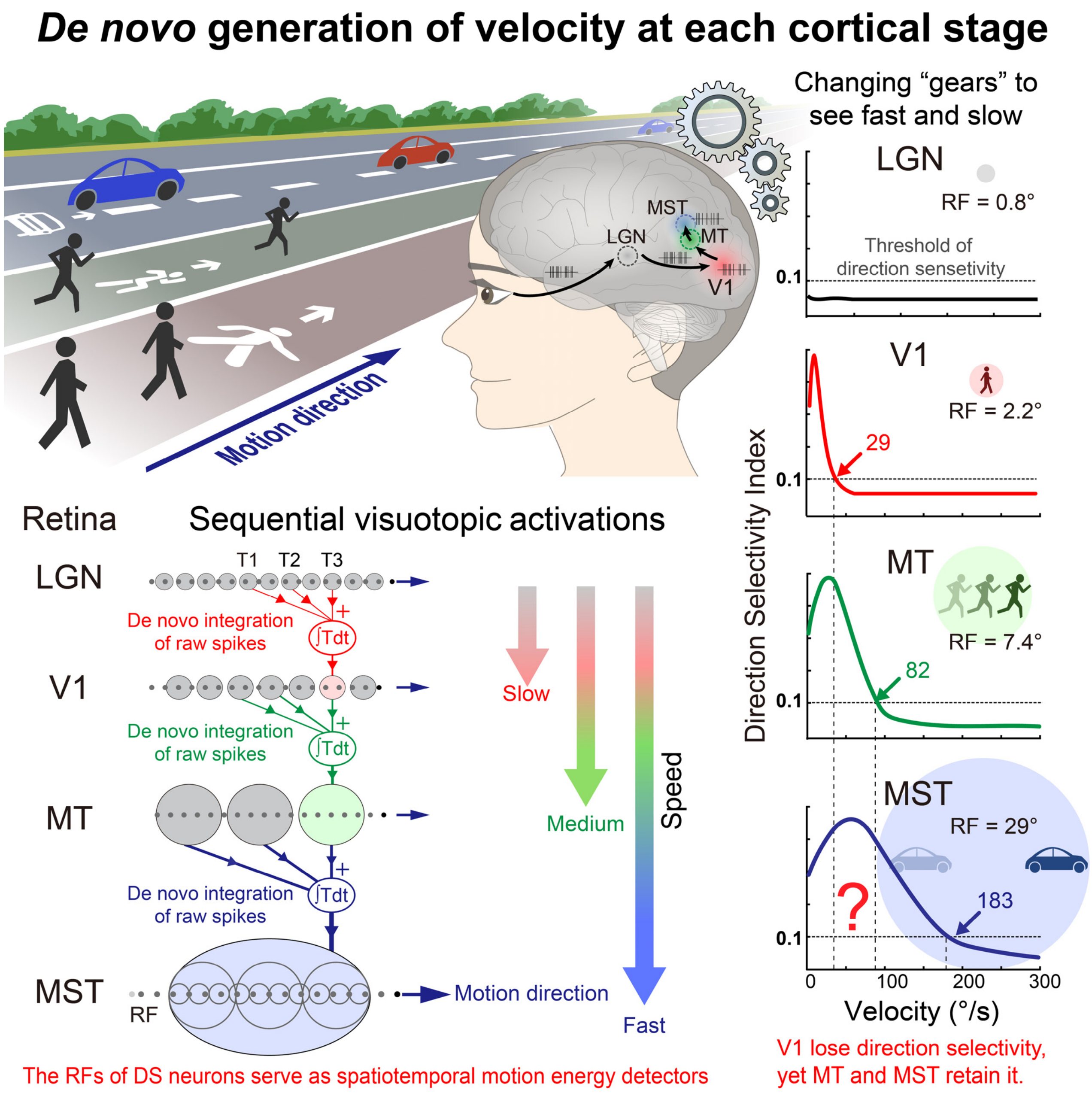


Graphic abstract shows how our hierarchically organized visual system employs each cortical area as a distinct "gear" to compute velocity anew for a wide spectrum of object speeds. The optimal and cutoff velocities of the tuning curves in V1, MT and MST are averaged velocities of recorded neurons across the three areas in awake macaques.

### Supplementary figures

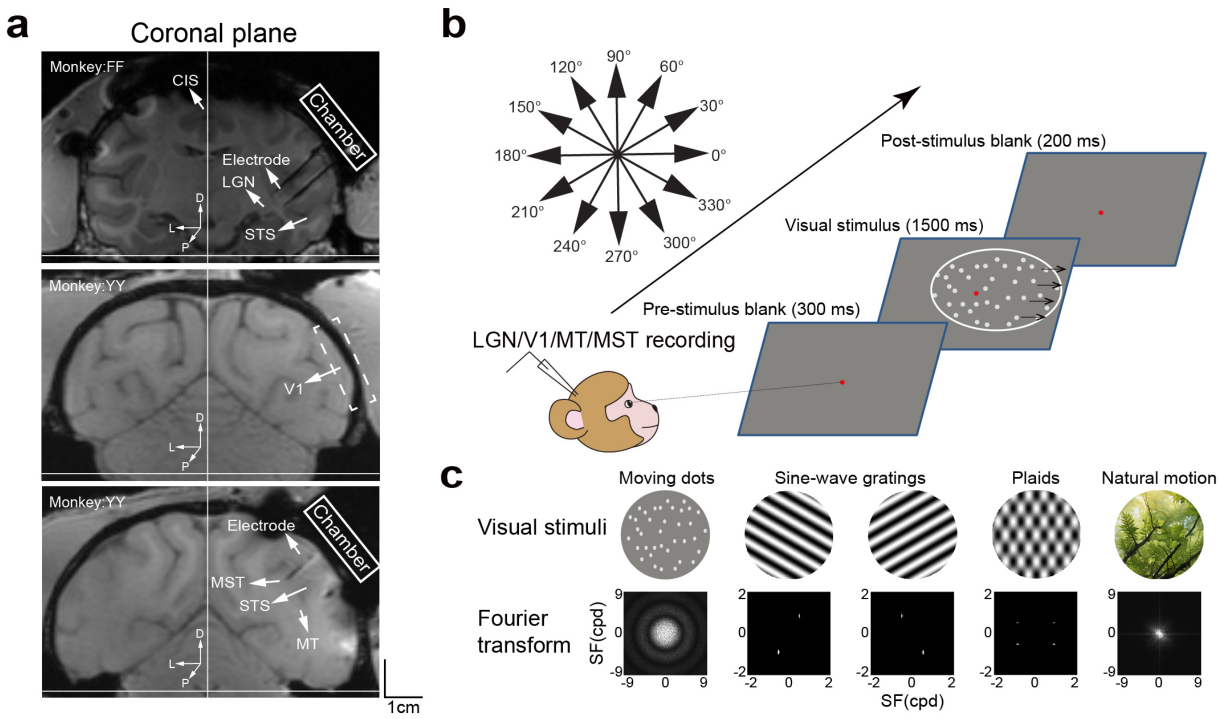


**Supplementary Fig.1 The experiment design, recording methods and visual stimuli. a.** MRI localization of visual brain areas of the lateral geniculate nucleus (LGN), the primary visual cortex (V1), the middle temporal area (MT) and the medial superior temporal area (MST) preparing for electrode penetrations. CIS, cingulate sulcus; STS, superior temporal sulcus. **b.** Stimulation settings and visual neuron recordings in LGN, V1, MT and MST along the hierarchy of the dorsal visual stream. **c.** The visual stimuli and their SF distributions after Fourier transformation.

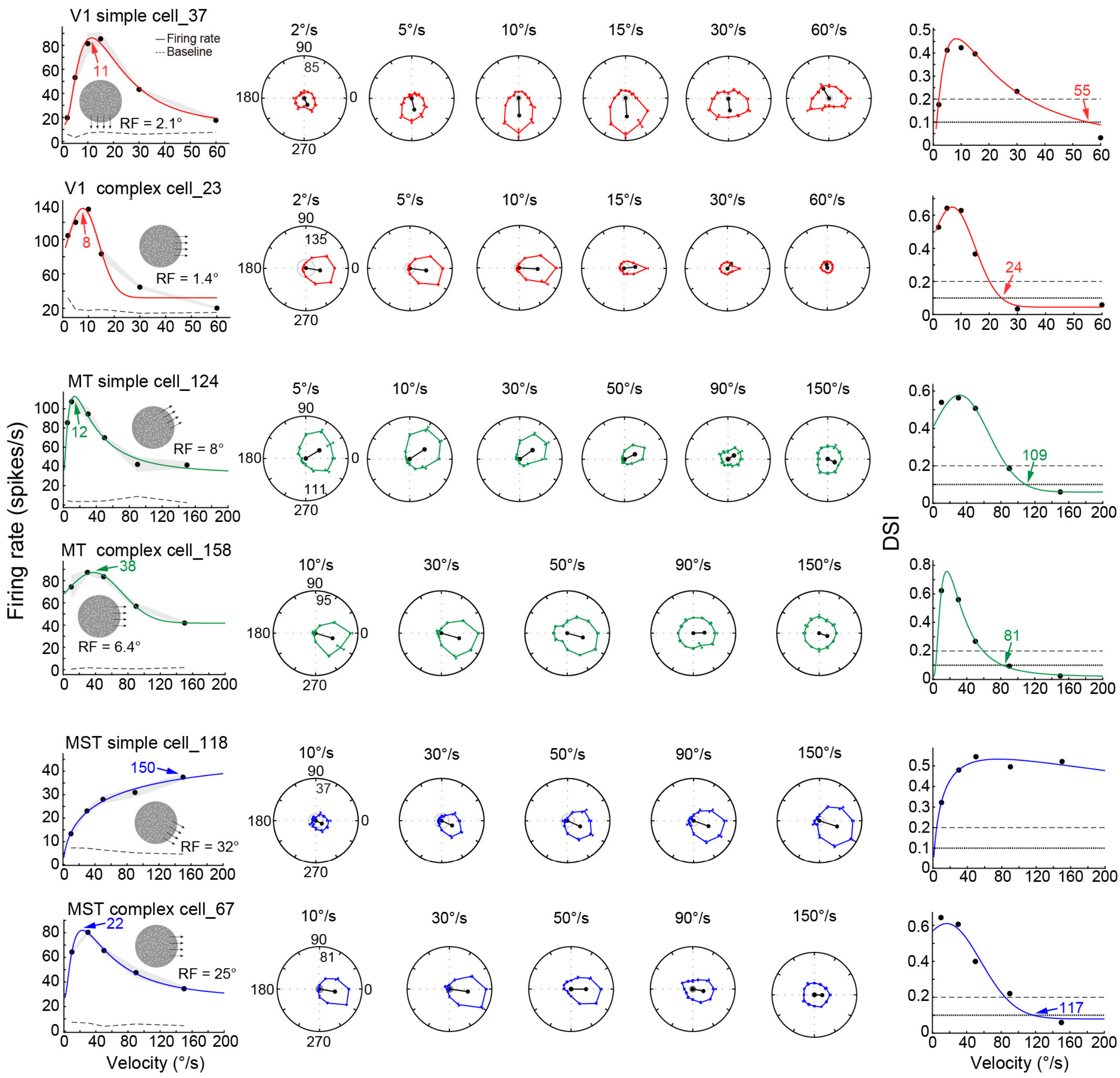


**Supplementary Fig.2 The velocity tuning examples of simple and complex cells in V1, MT and MST.** The classification of these examples is presented in **Fig.4a**. The left column shows the velocity tunings of each cell type across cortices, while the middle column exhibits the corresponding direction circular tuning curves of each cell type. The shade and error bar indicate the SEM, and also for the rest figures. The right column illustrates the DSI tuning curves of the corresponding cell types in the left and middle columns. The colored digits with arrow points indicate the optimal and cutoff velocities, respectively, and also for the rest figures.

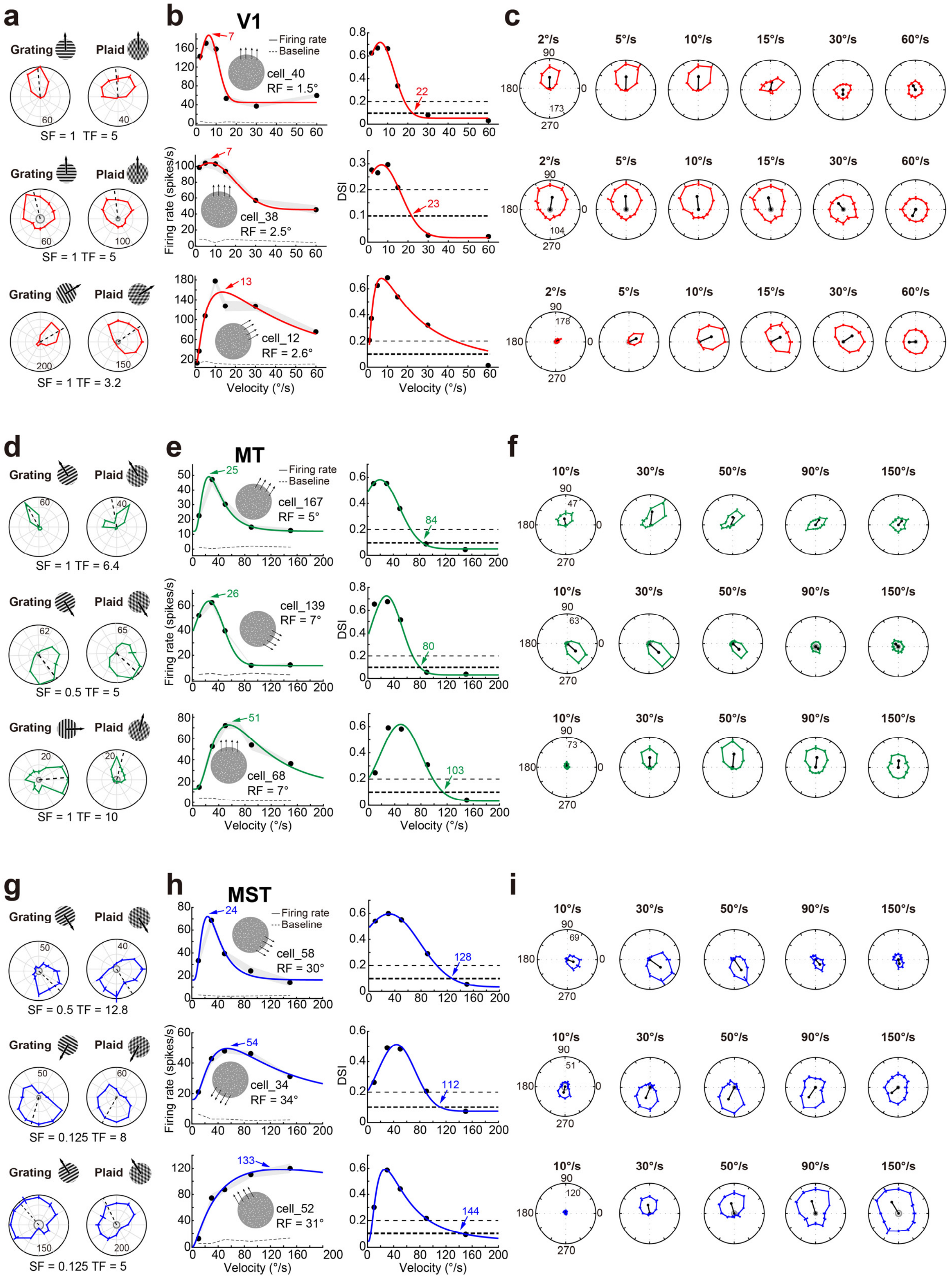


**Supplementary Fig.3 The component, pattern, and unclassified cell examples in V1, MT and MST.** **a.** The classification of three cell types in V1. **b.** The velocity tuning responses and DSI tuning curves for examples of the three cell types. **c.** The corresponding direction tuning curves in circular plots. **d.** The classification of three cell types in MT. **e.** The velocity tuning responses and DSI tuning curves for examples of the three cell types. **f.** The corresponding direction tuning curves in circular plots. **g.** The classification of three cell types in MST. **h.** The velocity tuning responses and DSI tuning curves for examples of the three cell types. **i.** The corresponding direction tuning curves in circular plots.

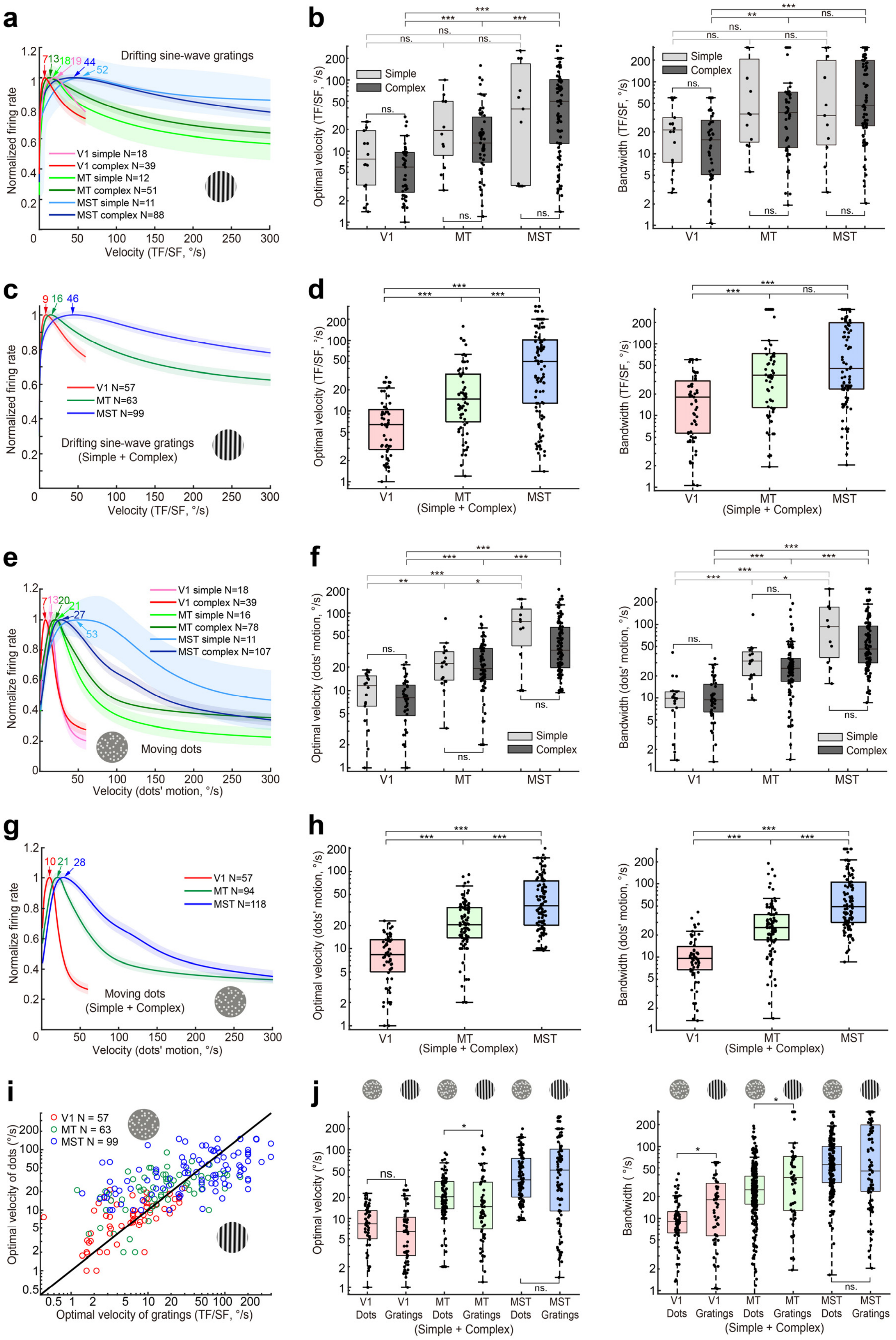


**Supplementary Fig.4 The velocity tuning responses of simple and complex cells to drifting sine-wave gratings and random dot motion across V1, MT and MST. a.** The velocity tuning responses of simple and complex cells to drifting gratings across cortices along the hierarchy. **b.** The comparison of optimal velocities and bandwidths for simple and complex cells within and across cortices of V1, MT and MST. The optimal velocity of simple cells: 11.51±2.10°/s, 32.30±8.65°/s and 78.33±28.45°/s for V1, MT and MST, respectively. The optimal velocity of complex cells: 7.15±1.08°/s, 23.93±4.29°/s and 74.86±8.31°/s for V1, MT and MST, respectively. The bandwidth of simple cells: 25.20±4.85°/s, 75.23±13.71°/s and 98.24±35.74°/s for V1, MT and MST, respectively. The bandwidth of complex cells: 20.47±2.92°/s, 75.23±13.71°/s and 108.51±111.60°/s for V1, MT and MST, respectively. **c.** The velocity tuning responses without separating simple and complex cells. **d.** The comparison of optimal velocities and bandwidths of the populations across V1, MT and MST. **e.** The velocity tuning responses of simple and complex cells to moving dots. **f.** The comparison of the optimal velocity and bandwidth of simple and complex cells within and across V1, MT and MST. The optimal velocity of simple cells: 10.36±1.33°/s, 25.45±4.70°/s and 77.19±14.86°/s for V1, MT and MST, respectively. The optimal velocity of complex cells: 8.62±0.90°/s, 24.94±1.95°/s and 49.01±3.91°/s for V1, MT and MST, respectively. The bandwidth of simple cells: 11.31±2.13°/s, 34.44±4.57°/s and 120.29±31.17°/s for V1, MT and MST, respectively. The bandwidth of complex cells: 11.83±1.26°/s, 33.04±3.63°/s and 75.45±6.98°/s for V1, MT and MST, respectively. **g.** The velocity tuning responses without separating simple and complex cells along the hierarchy. **h.** The comparison of the optimal velocity and bandwidth of populations across V1, MT and MST. **i.** The comparison of optimal velocities between moving fields of dots motion and drifting sine-wave gratings within each cortex along the hierarchy. **j.** The comparison of optimal velocities and bandwidths between the motion of dots and sine-wave gratings within each cortex of V1, MT and MST, respectively.

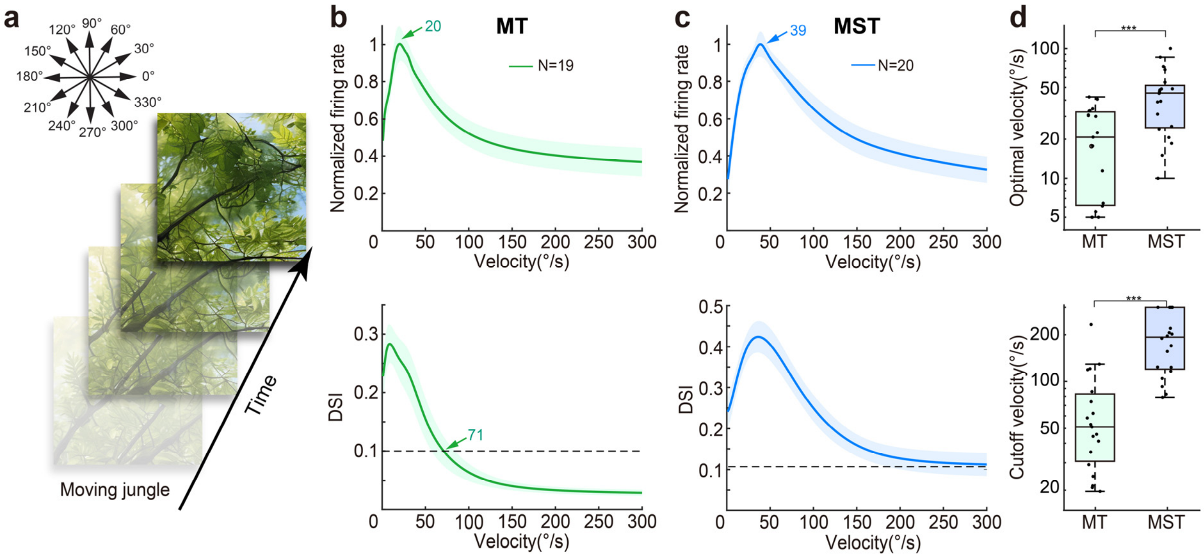


**Supplementary Fig.5 Natural stimuli for mapping MT and MST velocity and DSI tuning responses.** **a.** Example frames from a natural movie of a jungle scene. The velocity of natural movies used ranged from 5 to 200 °/s, and the movies presented within RFs of both MT and MST. **b.** The velocity and DSI tuning curves of MT DS neurons. **c.** The velocity and DSI tuning curves of MST DS neurons. **d.** The comparisons of optimal and cutoff velocities between MT and MST. Similarly to dots motion, both the optimal and cutoff velocities in MST are significantly higher than those in MT by using stimuli of natural image movies. The optimal velocity: 20.96±3.03°/s versus 44.33±5.29°/s for MT and MST, respectively. The cutoff velocity: 66.72±12.10°/s versus 178.53±15.10°/s for MT and MST, respectively.

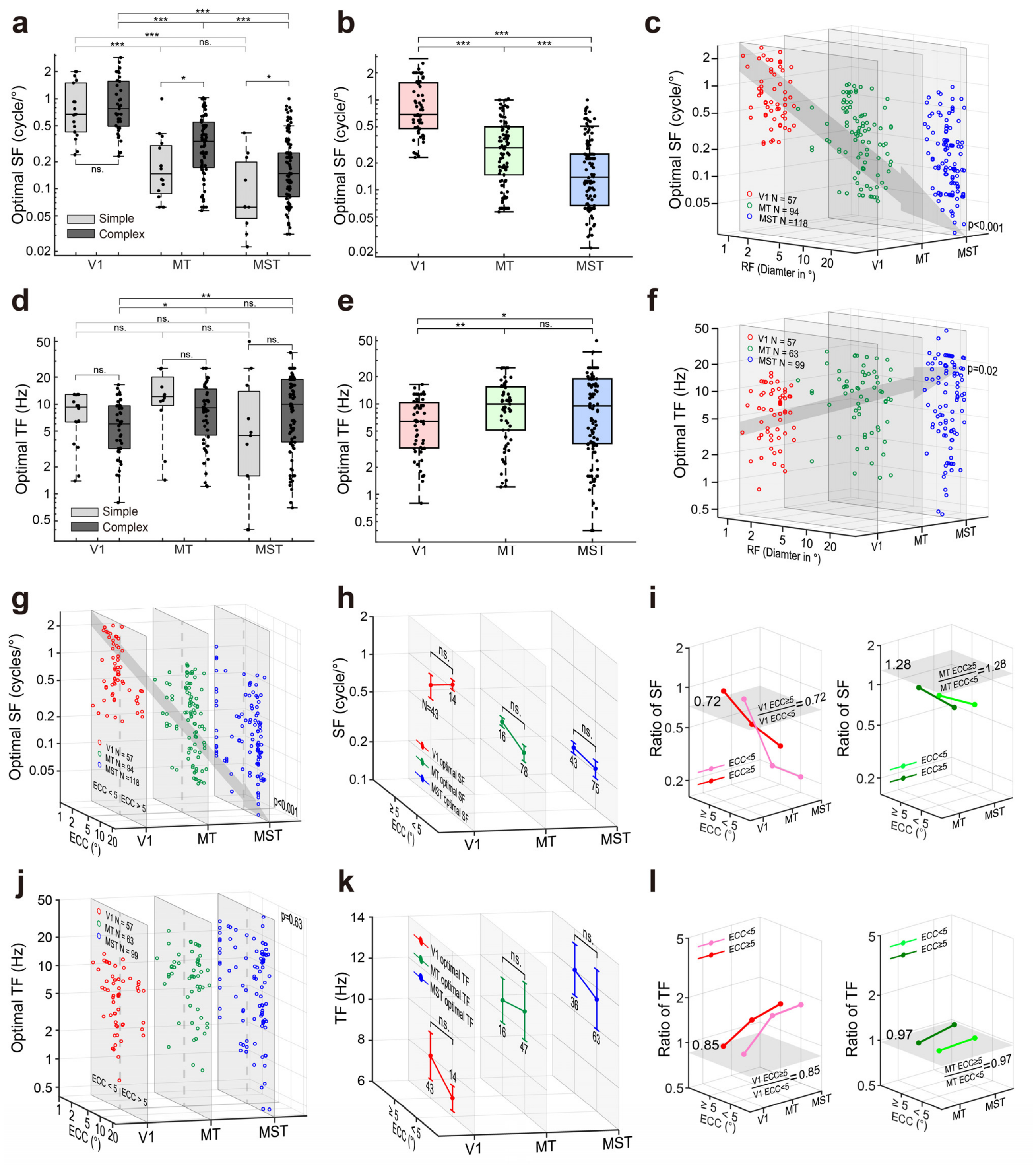


**Supplementary Fig.6 The comparison of optimal SFs and TFs cross cortices and eccentricities. a.** The comparison of optimal SFs for simple and complex cells across V1, MT and MST. There is no significant difference for V1 complex and simple cells in their optimal SF, consistent with previous reports(Foster et al., 1985). **b.** The comparison of optimal SFs without separating simple and complex cells. The optimal SF:0.99±0.09 cycle/°, 0.36±0.03 cycle/° and 0.22±0.02 cycle/° for V1, MT, and MST, respectively. **c.** The optimal SFs are reversely correlated with the increases of RFs across V1, MT and MST. **d.** The comparison of optimal TFs for simple and complex cells across V1, MT and MST. **e.** The comparison of optimal TFs without separating simple and complex cells across V1, MT and MST. The optimal TF: 7.23±0.54 Hz, 10.73±0.87 Hz and 11.8±0.96 Hz for V1, MT, and MST, respectively. **f.** The optimal TFs of DS neurons are weakly correlated with the increase of RF diameters across V1, MT and MST. **g.** The optimal SFs are reversely correlated with the increases of eccentricities across V1, MT and MST. **h.** The comparison of optimal SFs between below and above 5° eccentricities, respectively, across V1, MT and MST. **i.** The relative changes of the optimal SFs across cortices after normalizing to V1 and MT, respectively. The gray surfaces indicate the ratio levels of the optimal SFs for neurons above and below 5° eccentricities for V1 and MT, respectively. **j.** No correlation between optimal TFs and eccentricity across V1, MT and MST. **k.** The comparison of optimal TFs between below and above 5° eccentricities, respectively, across V1, MT and MST. **l.** The relative changes of the optimal TFs across cortices after normalizing them to V1 and MT, respectively. The gray surfaces indicate the ratio levels of the optimal TFs for neurons above and below 5° eccentricities within V1 and MT, respectively.

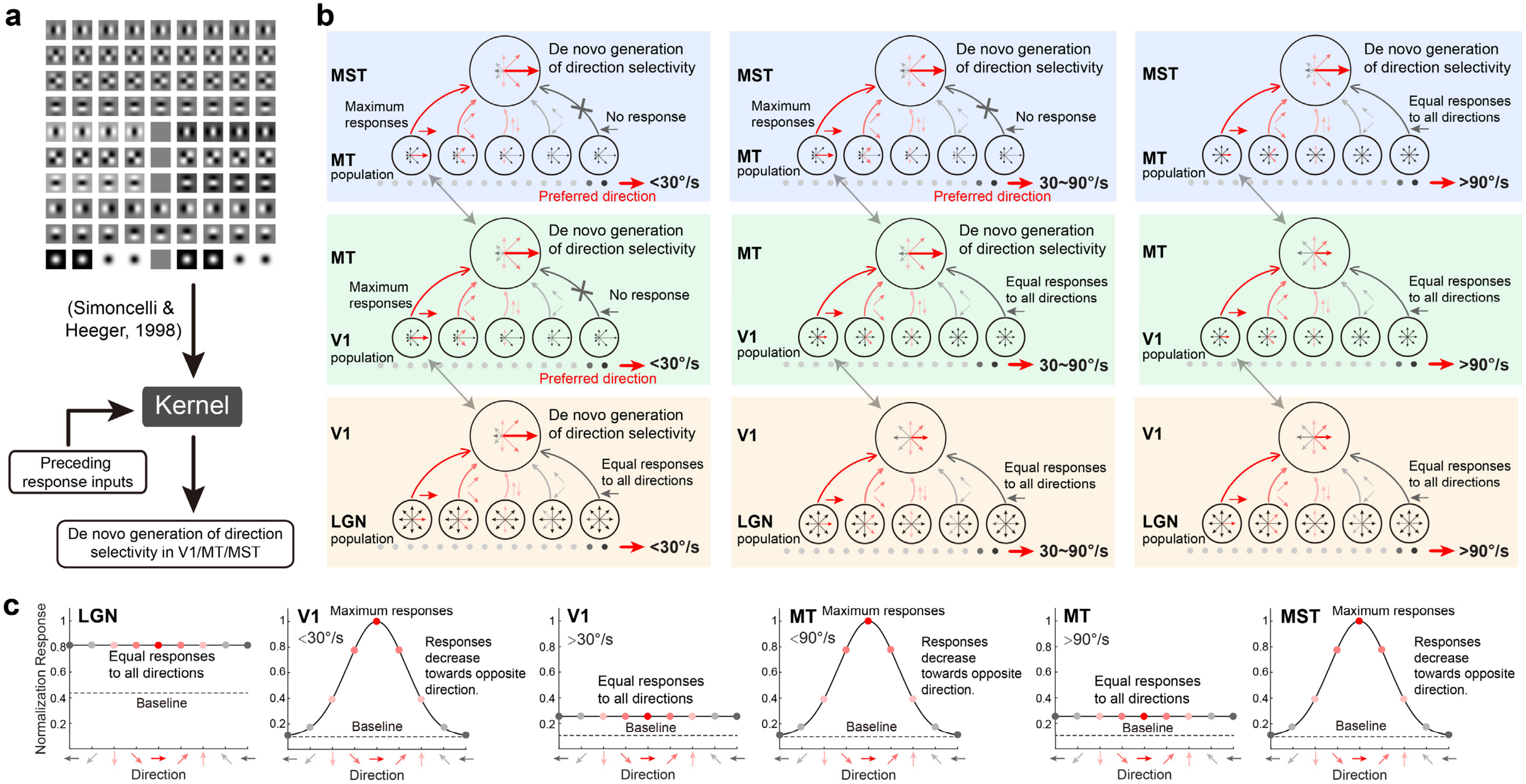


**Supplementary Fig.7 Schematic diagrams depict the cascaded spatiotemporal integration of presynaptic neuronal responses across cortices in the primate dorsal visual pathway. a. The mathematic filters (or kernels) replicate spatiotemporal RF substructures of V1 DS cells.** Ten basic spatiotemporal filters have been built to replicate the RF structures of DS cells(Freeman & Adelson, 1991; Simoncelli & Heeger, 1998), which are the core foundation of our cascaded spatiotemporal integration model. **b. The direction selective responses across V1, MT and MST associated with motion speeds.** For given moving stimuli, presynaptic spikes are maximum only when the stimulus is with the DS neuron's preferred motion direction. When the stimulus is away from the preferred direction, the firing rate decreases significantly. Direction selectivity varies across V1 and MT, respectively, which depending on different ranges of motion speeds.

**c. The direction selectivity depending on the motion speed, in V1, MT and MST, respectively.** LGN cells respond equally well to all motion directions, while V1 and MT alternatively lost their direction selectivity at their cutoff velocities of above 30 and 90°/s, respectively (**Fig.3**, please also see Mikami et al., 1986b; Churchland et al., 2005). The cutoff velocity of MST DS cells is beyond what we can possibly measure due to the fresh rate limitation of the stimulus monitor. Both the normalized responses and width of the tuning curves are just for illustration only.

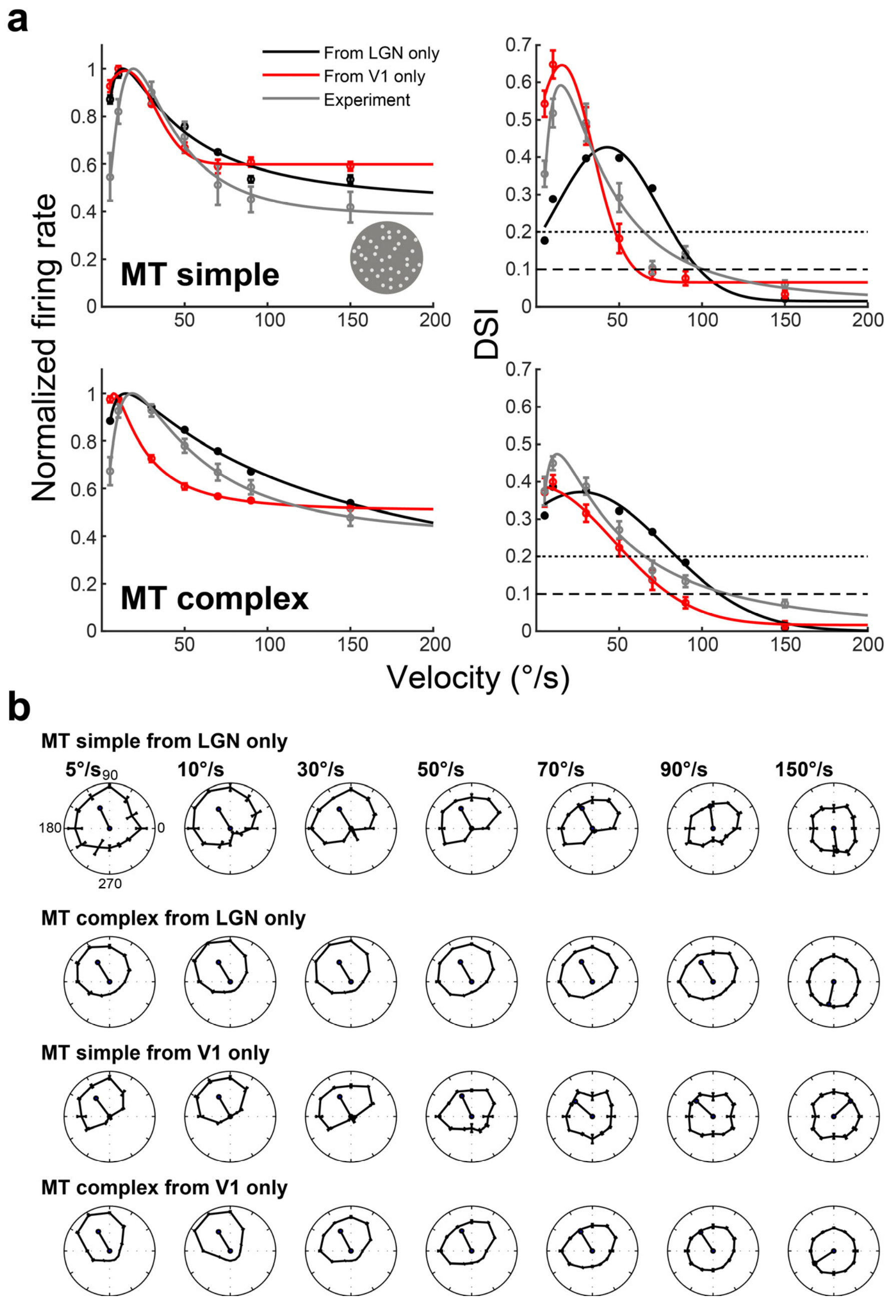


**Supplementary Fig.8 The model predicted results of velocity and DSI tuning responses in MT with and without V1 and LGN inputs, respectively. a.** The CSTI model simulated velocity and DSI tuning responses of MT simple and complex cells after blocking V1 and LGN inputs, respectively. **b.** The CSTI model simulated direction circular tuning curves at each speed for simple and complex cells in MT, after blocking LGN and V1 inputs, respectively.

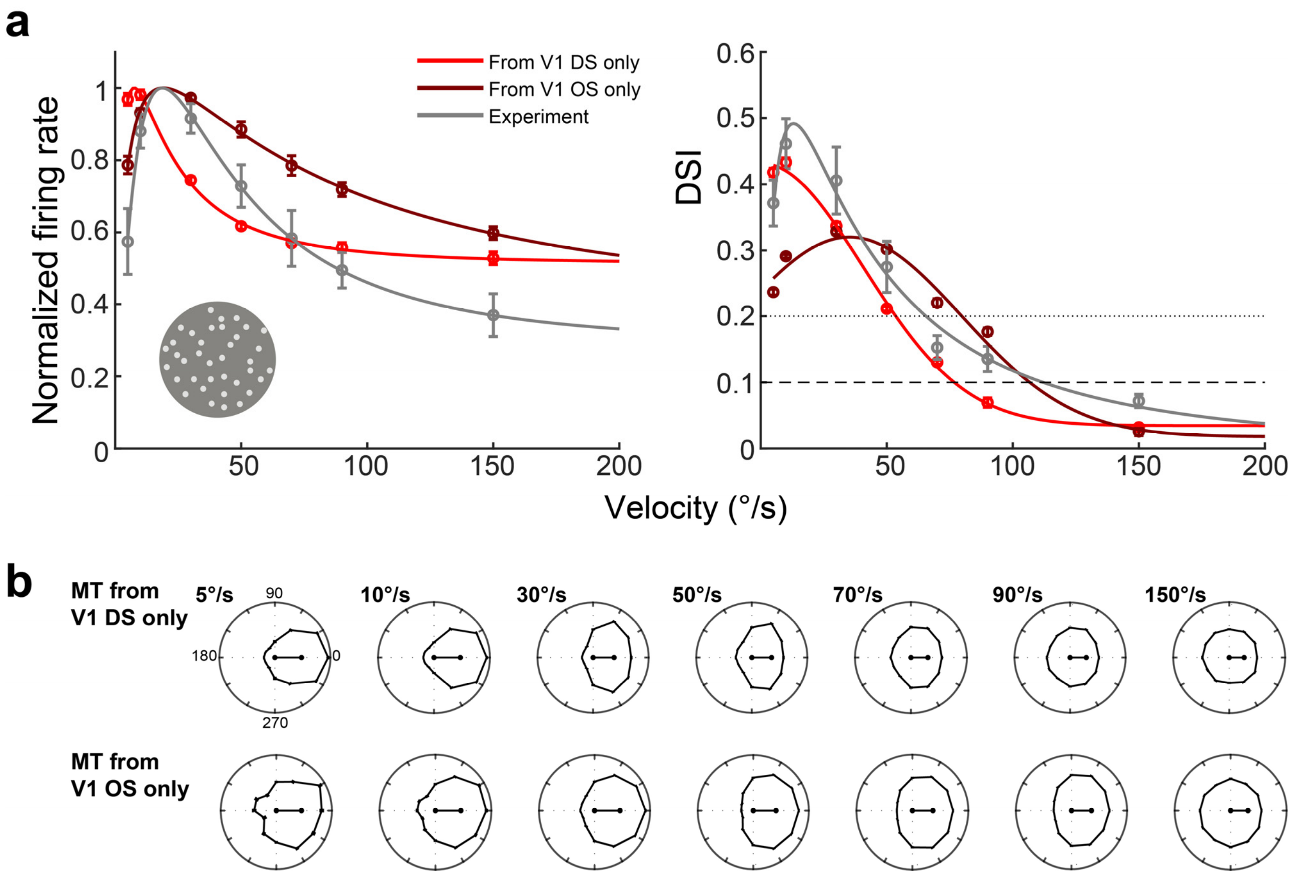


**Supplementary Fig.9 The model predicted results of velocity and DSI tuning responses in MT with only V1 DS and V1 OS inputs, respectively. a.** The CSTI model simulated velocity and DSI tuning responses of MT with V1 DS and V1 OS inputs only, respectively. **b.** The CSTI model simulated direction circular tuning curves at each speed for MT, from V1 DS and V1 OS inputs only, respectively. MT can generate its DS responses de novo irrespective of its presynaptic inputs.
